# A new phylogenetic framework for the genus *Kalanchoe* (Crassulaceae) and implications for infrageneric classification

**DOI:** 10.1101/2024.10.17.618389

**Authors:** Seraina E. Rodewald, David-Paul Klein, Ronen Shtein, Gideon F. Smith, Elizabeth Joyce, Diego F. Morales Briones, Sven Bernhard, Rokiman Letsara, Hannah Mertes, Philipp Hühn, Gudrun Kadereit

## Abstract

**Background and Aims:** *Kalanchoe* is a diverse genus in the Crassulaceae, with a centre of diversity in Madagascar and sub-Saharan Africa. The genus is known for its popularity in horticulture, its use as a model system for research on CAM photosynthesis and vegetative reproduction, its high invasive potential, and its use in traditional medicine. The genus-rank circumscription and infrageneric classification of *Kalanchoe* has been the subject of debate for centuries, especially regarding the status and rank of what is now treated as *K.* subg. *Bryophyllum* and *K.* subg. *Kitchingia*. We aim to generate a densely sampled phylogeny of *Kalanchoe s.l.* and evaluate the current infrageneric classification system.

**Methods:** We inferred a phylogenetic tree for *Kalanchoe* using a ddRAD sequencing approach, covering 70% of taxa and four out of five subgenera currently recognised in the genus.

**Key Results:** We recovered four well-supported clades, partially corresponding to the current subgeneric classification. *Kalanchoe* subg. *Calophygia* resolves as sister to the rest of the genus. The relationships among the three remaining clades, however, receive less support. The predominantly mainland African *K.* subg*. Kalanchoe* forms a strongly supported clade that resolves as sister to *K.* subg. *Bryophyllum*. These two clades are together sister to a clade containing mainly species from *K.* subg. *Kitchingia* and *K.* sect. *Pubescentes*.

**Conclusions:** The current subgeneric classification of *Kalanchoe* is partially backed up by our phylogenetic tree but requires further refinement. The tree topology suggests a Malagasy origin of the genus and one dispersal event to the African mainland, with subsequent dispersal from continental Africa to the Arabian Peninsula and Southeast Asia. The formation of bulbils on the leaf margin is restricted to a larger clade within *K.* subg. *Bryophyllum* and thus only evolved once. Our tree provides a framework for further taxonomic, evolutionary, and physiological research on the genus.

## Introduction

The genus *Kalanchoe* Adans. (Crassulaceae subfam. Cotyledonoideae; see Smith and Monro, submitted) comprises ca. 167 species (Smith and Figueiredo, 2023) (Fig.1) that are naturally distributed in Madagascar, Africa, the Arabian Peninsula, Southeast Asia, and northwest Australia. The genus is widely known for its cultivated species, such as *K. blossfeldiana* Poelln. and its hybrids and horticultural selections, which are worth millions of dollars to the horticultural industry each year (Van Voorst and Arends, 1982; Kahraman *et al*., 2022; Smith and Shtein, 2022). Some species of *Kalanchoe* are also used in traditional and homeopathic medicine for the pharmacological properties of the bufadienolides and flavonoids they contain (e.g. Nascimento *et al*., 2023; Assis De Andrade *et al*., 2023). The genus is additionally of considerable biological interest as a result of its diversity of vegetative reproduction modes (Smith *et al*., 2022), the most characteristic of which involves the formation of bulbils on leaf margins or inflorescences (Fig. 1 E, K) through complex developmental processes (Garcês *et al*., 2007; 2014). The pronounced ability of several *Kalanchoe* taxa to reproduce vegetatively contributes to the invasiveness of some species and nothospecies especially in places with a Mediterranean, subtropical, or tropical climate (Herrando-Moraira *et al*., 2020; Shtein and Smith, 2021; Gallo and Smith, 2024). Further, *Kalanchoe* species also have the ability to perform Crassulacean Acid Metabolism (CAM), a water-saving carbon concentration mechanism for photosynthesis. The expression of CAM varies within the genus, with some species having almost C_3_-like CAM behaviour, some showing strong inducible CAM, and some expressing strong, constitutive CAM (Kluge *et al*., 1991; 1993). As such, *Kalanchoe* has long been used as a model group for researching this photosynthetic pathway (e.g. Hartwell *et al*., 2016; Boxall *et al*., 2017; Yang *et al*., 2017; Winter, 2019). However, despite the great economic and biological importance of the genus, the phylogenetic relationships of *Kalanchoe* species remain unclear, and its infrageneric taxonomic classification has been the subject of debate for a long time.

**FIG. 1.**
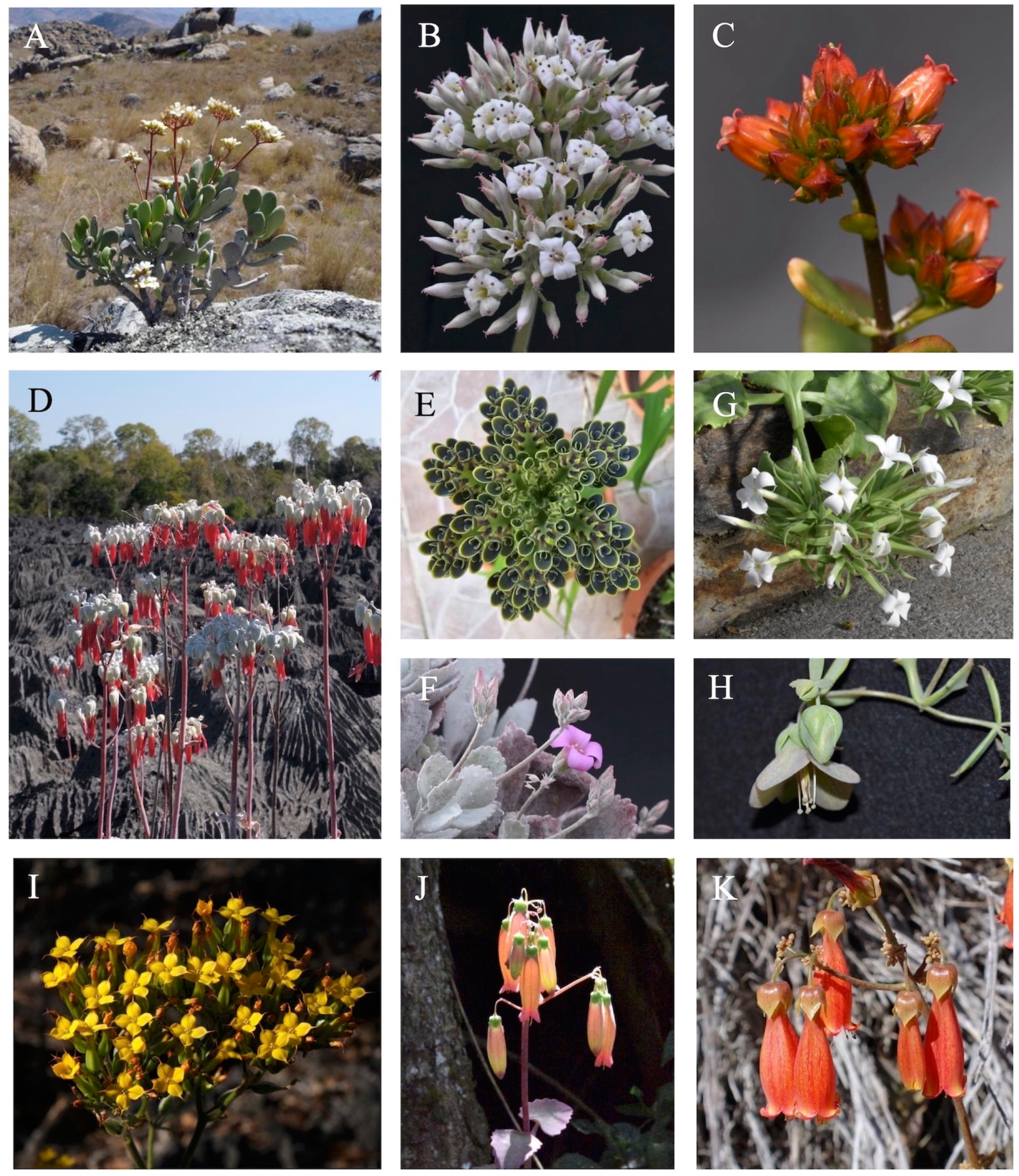
Morphological diversity in Kalanchoe A K. integrifolia var. integrifolia (K. subg. Calophygia) B K. synsepala (K. subg. Calophygia) C K. bracteata (K. subg. Calophygia) D K. bogneri (K. subg. Bryophyllum) E K. delagoensis (K. subg. Bryophyllum) F K. pumila (no subgeneric assignment) G K. schimperiana (K. subg. Kalanchoe) H K. beauverdii var. juelii (K. subg. Bryophyllum) I K. boisii (K. subg. Kalanchoe) J K. peltata var. stapfii (K. subg. Kitchingia) K K. miniata (no subgeneric assignment) Photographs: A, D, I, K David-Paul Klein, B, C, F – H, J Seraina E. Rodewald, E Ronen Shtein.

Since *Kalanchoe* was first described by Adanson (1763), names have been published for approximately 200 species, subspecies, and varieties in the genus. Morphologically, the genus is characterised, among others, by having tetramerous flowers with a (partially) fused calyx and corolla, an obdiplostemonous androecium and four apocarpous carpels, each with a nectar scale at the base. Over the years, several infrageneric treatments have been published in an effort to reflect species relationships, reviewed in detail by Smith (2023*g*; 2024*a*). Across most of the treatments, authors have divided *Kalanchoe* into three main taxa: *Kalanchoe s.s.*, *Bryophyllum*, and *Kitchingia*; however, the rank and status of *Bryophyllum* and *Kitchingi*a have been the subject of considerable debate. *Bryophyllum* Salisb. and *Kitchingia* Baker were initially described as separate genera by Salisbury (1805) and Baker (1881), respectively. Since then, *Bryophyllum* has mainly been treated as distinct from the rest of *Kalanchoe*, either at generic (e.g. Haworth, 1829; Baker, 1881; Baillon, 1885; Berger, 1930), subgeneric (e.g. Koorders, 1919; Maire, 1977; Descoings, 2006; Smith, 2022*a*), or section rank (Boiteau, 1947; Boiteau and Allorge-Boiteau, 1995; Descoings, 2003). *Kitchingia* has variously been treated as distinct from *Kalanchoe* and *Bryophyllum* at genus (Baker, 1881; Berger, 1930) or subgenus rank (e.g. Maire, 1977; Smith and Figueiredo, 2018; Smith *et al*., 2021*a*), as a section of *Kalanchoe* (e.g. Boiteau, 1947; Boiteau and Allorge-Boiteau, 1995), or as a synonym of *Kalanchoe* (e.g. Baillon, 1885; Hamet, 1907, 1908).

A recent comprehensive genus-wide taxonomic treatment was completed by Descoings (2006), who recognised three subgenera, i.e., the autonymic *K.* subg. *Kalanchoe*, *K.* subg. *Bryophyllum*, and the then newly described *K.* subg. *Calophygia* Desc. In the latter subgenus, Descoings (2006) included those species that he regarded as intermediate between *K.* subg. *Kalanchoe* and *K.* subg. *Bryophyllum*. However, this classification system of Descoings (2006) has not been adopted because of doubts about the monophyly of the subgenera that he recognised (see e.g. Smith and Figueiredo, 2018). Since the publication of Descoings (2006), extensive taxonomic work has been conducted at the specific, sectional, and subgeneric level (Smith and Figueiredo, 2018; Smith, 2020; 2021; 2022*c*; 2023*a*; *b*; *c*; *e*; *f*; 2024*b*; *d*; Shtein and Smith, 2021; Smith *et al*., 2021*a*; *b*). Following this recent taxonomic work, there are currently five subgenera in *Kalanchoe*: *Kalanchoe* subg. *Kalanchoe*, *K*. subg. *Bryophyllum* (Salisb.) Koord., *K.* subg. *Kitchingia* (Baker) Gideon F.Sm. & Figueiredo, *K.* subg. *Alatae* (Gideon F.Sm.) Raymond-Hamet ex Gideon F.Sm., and *K.* subg. *Calophygia* Desc. emend. Gideon F.Sm..

As these subgenera are currently circumscribed, the autonymic *K.* subg. *Kalanchoe* (Fig. 1 G, I) is the only one that is not endemic to Madagascar. Only a few species included in the subgenus are indigenous to Madagascar, while the majority of its representatives are distributed in continental Africa, with some found on the Arabian Peninsula and in Southeast Asia (Smith *et al*., 2019) reaching into northwest Australia (Kenneally, 1983). All other subgenera are endemic to Madagascar but especially *K.* subg. *Bryophyllum* (Fig. 1 D, E, H) contains several species that have become naturalised and even invasive in large areas of mild climatic zones (Shtein and Smith, 2021; Smith, 2023*d*). While *K.* subg. *Bryophyllum* were published more than 100 years ago (Koorders, 1919), which established the autonymic *K.* subg. *Kalanchoe*, a validly published combination for *K.* subg. *Kitchingia* (Fig. 1 J) became available much later (Smith and Figueiredo, 2018) and was more recently amended to contain only the two species *K. gracilipes* Baill. and *K. peltata* Baill. (Smith *et al*., 2021*a*). *Kalanchoe* subg. *Alatae* was established for the two epiphytic species *K. uniflora* (Stapf) Raym.-Hamet and *K. porphyrocalyx* (Baker) Baill. (Smith, 2023*a*). Finally, *K.* subg. *Calophygia* (Fig. 1 A–C) was reinstated by Smith (2023*f*) and amended substantially from the original definition *sensu* Descoings (2006), now exclusively containing the Malagasy ‘woody clade’ that had been consistently treated under *Kalanchoe s.s.*, i.e., *K.* subg. *Kalanchoe*, throughout most of the taxonomic history of the genus (Smith, 2023*f*).

As currently circumscribed, the subgenera recognised in *Kalanchoe* are distinguished based on a suite of flower, life form, and vegetative reproductive characters, and to a lesser extent on their natural geographical distribution ranges.

One of the main characters initially used to distinguish *Bryophyllum* from *Kalanchoe s.s.* was the morphology of the calyx, where *Bryophyllum* was described as having a large and substantially fused, often flimsy calyx with short lobes, while *Kalanchoe* was defined as having (almost entirely) free sepals with at most a very short tube (Salisbury, 1805; de Candolle, 1828; Hamet, 1907; Descoings, 2006). The later description of intermediate species sparked considerable debate on the usefulness of calyx characters and, as a consequence, the desirability of distinguishing the two taxa as separate genera (see e.g. de Candolle, 1828; Dalzell, 1852; Hamet, 1907; 1908; Descoings, 2006). At present the degree of fusion of the sepals is again, partly, regarded as informative to characterise *Bryophyllum* but now at the rank of subgenus (Smith, 2022*a*; 2024*d*). When *Kitchingia* was described as a genus some 80 years after the genus name *Bryophyllum* was published, the calyx was described as being small with the tube and lobes being of similar lengths (Baker, 1881), thus showing a somewhat intermediate character state between *Kalanchoe* and *Bryophyllum*. *Kalanchoe* subg. *Alatae* was defined as having a flattened calyx tube that is shorter than the free sepal segments (Smith *et al*., 2021*b*) and *K.* subg. *Calophygia* is characterised by having a “prominent, sometimes distinctly succulent” calyx with the sepals being either free or fused (Smith, 2023*f*).

To distinguish among the subgenera the shape of the corolla tube was interpreted as follows: in *K.* subg. *Kalanchoe*, it is enlarged towards the middle or more often at the base and often constricted below the petal lobes; in *K.* subg. *Bryophyllum* the corolla tube is straight or slightly widening, often constricted above the ovaries; in *K.* subg. *Kitchingia* the corolla tube is tubular to campanulate and not constricted; in *K.* subg. *Alatae* the corolla tube is inflated in the lower half, and in *K.* subg. *Calophygia* it is “±quadrangular-urceolate, tapering to [the] mouth” (see e.g. Berger, 1930; Descoings, 2006; Smith *et al*., 2021*a*; *b*; Smith, 2023*f*).

Another important, albeit debatable, character to distinguish the infrageneric taxa is the point of insertion of the filaments in relation to the corolla tube. For *Bryophyllum*, across treatments, regardless of the taxonomic rank assigned to it, the filaments have been described as being inserted below the middle. For *Kalanchoe s.s.*, however, the position of filament insertion was interpreted by Berger (1930) as being variable, while Descoings (2006), in his more narrowly defined circumscription of the taxon (as the autonymic *K.* subg. *Kalanchoe*), described the filaments as normally being inserted above the middle of the corolla tube, often towards the top but sometimes towards the middle. Smith and Figueiredo (2018) described the filaments of *K.* subg. *Kalanchoe* to be inserted “± medially” and in the most recent treatment of the subgenus (Smith, 2024*d*), the filaments are described to be “± inserted in [the] upper ½ of [the] corolla tube”. For *Kitchingia*, in the original description as well as in the current classification of the taxon, the filaments are described as being inserted above the middle. However, the broader definitions of the taxon by both Baillon (1885) and Smith and Figueiredo (2018) also included species with filaments inserted below the middle of the corolla tube, and thus in the diagnosis of *K.* subg. *Kitchingia sensu* Smith and Figueiredo (2018) they were described to be “ ± inserted in [the] lower third of [the] corolla tube”. For *K.* subg. *Alatae*, the filaments are described as being inserted in the lower half of the corolla tube and for *K.* subg. *Calophygia* as being inserted in the middle or above the middle of the corolla tube.

Other characters deemed as important in distinguishing the major infrageneric taxa in *Kalanchoe* include the length of the style relative to the ovary, divergence of the carpels, length of the nectar scales, and flower orientation. The length of the style relative to the ovary is used as a distinguishing character for the description of all subgenera except *K.* subg. *Calophygia*: the style being shorter than the ovaries or rarely equal in length in *K.* subg. *Kalanchoe* (Berger, 1930; Descoings, 2006), similar in length in *K.* subg. *Alatae* (Smith *et al*., 2021*b*), but longer than the ovaries in *K.* subg. *Bryophyllum* (Smith, 2022*a*) and *K.* subg. *Kitchingia* (Smith *et al*., 2021*a*).

One of the most important characters used to distinguish *K.* subg. *Kitchingia* from the other subgenera are diverging carpels present in *Kitchingia s.s.* (Baker, 1881; Smith, *et al*. 2021*a*), while again, in the wider definitions of the taxon *sensu* Baillon (1885) and *sensu* Smith and Figueiredo (2018), the character is variable. The nectar scales were described as being variable in length and width in *K.* subg. *Kalanchoe* (Berger, 1930) but shorter than wide or similar in length and width but never linear for both *K.* subg. *Bryophyllum* (Smith, 2022*a*) and *K.* subg. *Kitchingia* (Smith *et al*., 2021*a*), linear in *K.* subg. *Alatae* and “generally wider than long” in *K.* subg. *Calophygia*. Furthermore, flowers in *K.* subg. *Bryophyllum*, *K.* subg. *Kitchingia*, and *K.* subg. *Alatae* are pendulous (see e.g. Smith *et al*., 2021*a*; *b*; Smith, 2022*a*) while they are erect or spreading in *K.* subg. *Kalanchoe* and *K.* subg. *Calophygia* (see e.g. Smith and Figueiredo, 2018; Smith, 2023*f*).

In terms of vegetative reproduction the development of bulbils on the leaf margin (Fig. 1 E), i.e. being “phyllo-bulbiliferous” (see Shtein and Smith, 2021), is a diagnostic feature of *Bryophyllum*, regardless of the rank at which it is treated (Smith, 2022*a*). However, not all species currently accepted in *Bryophyllum* show this trait.

Whether these traits overall contribute to the demarcation of clades in *Kalanchoe s.l.* remains to be tested in a phylogenetic context.

In recent taxonomic work in *Kalanchoe*, the subgenera have often been treated individually, with a comprehensive genus-wide classification not being available to date. Some species are therefore currently not unambiguously assigned to a subgenus. For example, species excluded from *K.* subg. *Calophygia* by Smith (2023*f*) were mostly placed in one of the other subgenera in Smith (2024*a*), while *K. aromatica* H.Perrier, *K. bouvetii* Raym.-Hamet & H.Perrier, *K. pumila* Baker, *K. chapototii* Raym.-Hamet & H.Perrier, *K. quadrangularis* Desc., and *K. tuberosa* H.Perrier, as well as *K.* sect. *Pubescentes* (A. Berger) Gideon F.Sm., have thus far not been placed. In addition, the sectional classification of *Kalanchoe* is still to be completed. Eleven sections have been published recently (Smith, 2020; 2022*c*; 2023*b*; *c*; *e*; 2024*b*; *c*; *d*) but most species (especially in *K.* subg. *Kalanchoe*) have not been classified into sections. As a consequence, the sections have not been discussed in relation to each other, which make a comparison among them difficult. A comprehensive, genus-wide phylogeny will enable the testing of the validity of existing subgeneric and sectional concepts.

Thus far there is insufficient molecular evidence to test the current morphological infrageneric classification of *Kalanchoe.* The most detailed molecular phylogenetic study of *Kalanchoe* published to date is that of Gehrig *et al*. (2001), who generated a phylogenetic tree of 54 species, about ⅓ of the species recognised in the genus, based on one nuclear marker (ITS). They found that species from *Bryophyllum* and *Kitchingia* fell inside *Kalanchoe*, and consequently proposed three sections—‘Kitchingia’, ‘Bryophyllum’ and ‘Eukalanchoe’, i.e., *Kalanchoe*. However, because of the limited amount of data included, the relationships of the major clades were retrieved as a large polytomy, rendering their relationships unclear. As such, the classification suggested by Gehrig *et al*. (2001) remains to be validated with additional data. Since the study of Gehrig *et al*. (2001), there has been no comprehensive updated phylogenetic framework for the genus. The phylogenetic relationships among the genera included in Crassulaceae subfam. Cotyledonoideae as amended by Smith and Monro (submitted) have been studied extensively (Mort *et al*., 2001; Nowell, 2008; Messerschmid *et al*., 2020; Liu *et al*., 2023; Han *et al*., 2024), and confirm the monophyly of the genus *Kalanchoe*; however, only a small number of *Kalanchoe* species were included in these family- or subfamily-wide studies. As such, a rigorous phylogenetic analysis of *Kalanchoe* with more extensive sampling and more data is needed to test the current taxonomic classification and to provide a framework for future evolutionary studies in the group.

The aim of this study is to generate a comprehensive phylogenetic tree of the genus *Kalanchoe* using a modified double digest Restriction-site Associated DNA sequencing (ddRADseq) approach that targets long loci. This approach has been shown to be effective in other Crassulacean lineages (Hühn *et al*., 2022; Messerschmid *et al*., 2023). Based on this new phylogeny, we aim to evaluate if the current infrageneric classification is supported by molecular evidence. The updated phylogeny will provide a framework for further evolutionary, horticultural, physiological, and taxonomic research on the genus. Future phylogenetically informed studies will improve our understanding of the evolution of traits such as CAM photosynthesis, vegetative reproduction, flower morphology, and growth form.

## Material and Methods

### Sampling and DNA extraction

We sampled 184 accessions of *Kalanchoe* belonging to 138 taxa, covering approximately 70% of the species diversity and four out of the five currently accepted subgenera (Table S1). *Kalanchoe* subg. *Alatae*, which comprises the two species *K. porphyrocalyx* and *K. uniflora,* was not included due to the unavailability of material. We included four outgroup species of the genera *Adromischus* Lem., *Cotyledon* L., and *Tylecodon* Toelken, which, together with *Kalanchoe*, make up Crassulaceae subfam. Cotyledonoideae A.Berger emend. Gideon F.Sm. (Smith and Monro, submitted). We obtained fresh leaf samples from the living collection of Botanischer Garten München-Nymphenburg (M) and Botanischer Garten der Johannes Gutenberg-Universität Mainz (MJG) that are vouchered at the Botanische Staatssammlung München (M) or silica-dried material received from The Yehuda Naftali Botanic Garden of Tel Aviv University (TELA), Botanischer Garten und Botanisches Museum Berlin-Dahlem (B), the Muséum national d’Histoire naturelle in Paris and the private living collection of one of us (GFS). The latter accessions were collected between 1981 and 2001 and are vouchered at the H.G.W.J. Schweickerdt Herbarium (PRU). Additional silica material samples were collected in the field in Madagascar in October 2023, and one sample was obtained from Conservatoire et Jardin botaniques de la Ville de Genève, collected in the field in February 2022.

Prior to DNA extraction, we lyophilised fresh material to increase DNA yield. We extracted DNA with the DNEasy Plant Mini-Kit (QIAGEN, Venlo, Netherlands) following the manufacturer’s instructions. DNA quantity was measured with a Qubit 3.0 Fluorometer (Thermo Fisher Scientific, Waltham, MA, USA), and DNA quality was determined with agarose gel electrophoresis.

### Library preparation and sequencing

We followed the protocol of Hühn *et al*. (2022) to prepare libraries for ddRAD sequencing. This protocol targets long loci (300–600 bp) and has previously been successfully used for a coalescent-based approach in Crassulaceae (Hühn *et al*., 2022). We estimated that double DNA digestion with the restriction endonucleases (REase/s) *BamHI* (restriction site: G’GATCC) and *KpnI* (restriction site: GGTAC’C) would result in ca. 5000 300–600 bp DNA fragments in *Kalanchoe* with an *in silico* digestion using the CLC genomics workbench v9.5.5 (QIAGEN, Venlo, Netherlands) on the genome of *K. fedtschenkoi* Raym.-Hamet & H.Perrier (BioProject PRJNA397334). Based on these results, we used these REase/s to digest 200ng genomic DNA per sample.

After digestion, adapters containing an 8 bp unique barcode were ligated to the DNA fragments. Following Hühn *et al*. (2022), we used an adapter design where all possible fragment types can be sequenced, including fragments with the same restriction motif on both ends. This can be achieved by using both a common adapter and an adapter containing a unique barcode for both restriction motifs. The reaction was subsequently cleaned using magnetic beads. To avoid index hopping (Van Der Valk *et al*., 2020), we added an additional indexing step to the protocol of Hühn *et al*. (2022). In this step, a 12-cycle 3-step PCR was performed, where each sample received a unique combination of index primers. The samples were then multiplexed, and the libraries were cleaned with NucleoSpin Gel and PCR Clean-up columns (Macherey-Nagel, Düren, Germany). For size selection, we used a Blue Pippin (Sage Science, Beverly, MA, USA) to select fragments of 440-810 bp length, including adapters and primers (i.e., a length of 300-670 bp of targeted insert DNA fragments). To remove potential heteroduplexes, we performed a one-cycle reconditioning PCR. The PCR reaction was cleaned with columns. For the final purification with magnetic beads (NucleoMag NGS kit, Macherey-Nagel, Düren, Germany), we used a ratio of 0.65:1 bead solution to the library to discard small fragments originating from heteroduplexes that had passed the size selection. The fragment length distribution in the final library was assessed with a TapeStation automated electrophoresis (Agilent, Santa Clara, CA, USA). The libraries were subsequently sequenced on an Illumina MiSeq 300 bp paired-end at the Genomics Service Unit of the Faculty of Biology of Ludwig-Maximilians-Universität München.

### Data assembly

For trimming adapters from the demultiplexed raw reads, we used Cutadapt v3.5 (Martin, 2011). Sequences with wrongly assigned barcodes were discarded. We used an error threshold of 12.5% at the 3’-end instead of the default 10% to allow adapters to be removed with a sequencing error of one in eight bases to ensure that short fragments did not contain adapter sequences that could influence downstream analysis. Read quality was assessed with FastQC v 0.11.9 (Andrews, 2010) and MultiQC (Ewels *et al*., 2016) before and after trimming.

For the *de novo* assembly of our reads, we used ipyrad v 0.9.90 (Eaton and Overcast, 2020). We set ‘max_alleles_consens’ (parameter 18) to four, allowing up to four alleles per locus and excluding any locus with more alleles. This was done to account for the high variation in ploidy levels in *Kalanchoe*—ploidy varies from diploid to decaploid across the genus (Smith, 2022*b*), but remains unknown for many species. We set ‘max_Indels_locus’ (parameter 23) to 24, allowing for up to 24 indels per locus and accounting for our elevated read length compared to standard RADseq reads. ‘max_shared_Hs_locus’ (parameter 24) was set to 0.7 following (Hühn *et al*., 2022). To choose an appropriate clustering threshold for our data (parameter #14), we performed separate test series of ipyrad assemblies for within-sample clustering and between-sample clustering. To determine the optimal within-sample clustering threshold, we ran steps 1-5 of the ipyrad pipeline with the clustering threshold set to values from 0.80–0.98 with an increment of 0.02 and evaluated the following assembly metrics: the total number of clusters, the average depth, the number of loci filtered by maximum heterozygosity, and heterozygosity (Supplementary Fig. S1a). With an increasing clustering threshold, the number of clusters increased steadily while the average depth decreased. The heterozygosity and number of loci filtered by maximum heterozygosity decreased steeply above a clustering threshold of 0.94, which indicates a balance between over-merging and under-merging at this threshold (see Hühn *et al*., 2022). We thus chose 0.94 as the optimal clustering threshold for within-sample clustering. To select the between-sample clustering threshold, we ran steps 6–7 of the ipyrad assembly pipeline with the clustering threshold set to values from 0.80–0.98 with an increment of 0.02. As input, we used the assembly generated until step 5, with a within-sample clustering threshold of 0.94. We plotted and evaluated the number of retained loci, the total number of variable sites, the average number of variable sites per locus, and the percentage of missing data (Supplementary Fig. S1b). The percentage of missing data was around 96% over all clustering thresholds and did not increase with an increasing clustering threshold because loci covering less than four samples were filtered out following the ipyrad default settings. We thus did not further consider the percentage of missing data to inform our choice of between-sample clustering threshold. The number of retained clusters and the total number of variable sites peaked at a clustering threshold of 0.94 while the average number of variable sites per locus dropped significantly after a clustering threshold of 0.90. We thus chose a between-sample clustering threshold of 0.90.

### Phylogenetic inference

We used the concatenated alignment including all loci from the ipyrad assembly to infer a maximum likelihood tree using IQ-TREE v2.2.2.7 (Minh *et al*., 2020) under the GTR +F +G4 substitution model. We hereafter refer to this phylogenetic tree as the ‘concatenated tree’. Node support was evaluated using 1000 ultra-fast bootstrap replicates (UFBS) (Hoang *et al*., 2018).

We also estimated a species tree using a two-step coalescent approach, hereafter called the ‘coalescent tree’. For this tree, we used a filtered dataset because multi-species coalescence summary methods such as ASTRAL are very sensitive to gene tree estimation error and, thus, to missing data (Molloy and Warnow, 2018). Loci were parsed and extracted using scripts from Hühn *et al*. (2022). We selected loci with a minimum of 100 bp that contain at least 10% of samples and have a minimum of ten parsimony informative sites. Gene trees were subsequently estimated using IQ-TREE under the GTR + F + G4 model and branches with less than 70% UFBS support were collapsed. These gene trees were then used to estimate the species tree under the coalescent process using ASTRAL v1.16.3.4 (Zhang and Mirarab, 2022) with default settings. Branch support for the coalescent tree was estimated as local posterior probabilities (PP) (Sayyari and Mirarab, 2016).

To assess the robustness of both the concatenated and the coalescent tree and explore possible conflict, we used Quartet Sampling (QS; Pease *et al*., 2018). In this method, all samples of a phylogeny are split into four non-overlapping subsets at each internal branch. From each of the subsets, taxa are randomly selected to form quartets for which quartet phylogenies are calculated that can be either concordant or discordant with the tree or be uninformative. With this, the method aims to differentiate between conflict and lack of support in phylogenetic inferences. We ran QS on both the concatenated and coalescent trees using the concatenated alignment of Single Nucleotide Polymorphisms (SNPs) with IQ-TREE as the tree inference engine (Nguyen *et al*., 2015). The minimum overlap for a quartet to be considered was set to 100 SNPs, corresponding to about four loci. Quartet Sampling outputs three scores at each internal branch: (1) the quartet concordance (QC) score, which is a measure for the number of concordant versus discordant quartets at this bipartition, (2) the quartet differential (QD), which shows the relative frequency of the two possible discordant topologies, and (3) the quartet informativeness (QI), which is the proportion of quartets that were informative, i.e. that passed the likelihood cutoff. We used the default likelihood cutoff of two; i.e., the most likely quartet topology needs to be at least twice as likely for the quartet to be counted as concordant or discordant. For the tips, quartet sampling outputs the quartet fidelity (QF), which is the proportion of concordant quartets inferred that included this sample.

## Results

### Data recovery and assembly

We obtained an average of 383,774 (70,569–970,836) reads per sample from Illumina paired-end sequencing with an average sequence length of 263 bp (178–291 bp) after adapter trimming.

The ipyrad assembly resulted in a final number of 28,927 loci, with an average sample coverage of 1,158 loci per sample for the ingroup and 19 for the outgroup. The average locus length is 324 bp, with an average of 25 SNPs and 7 samples per locus. The number of shared loci between samples shows a strong clustering into clades, see supplementary Fig. S2.

The filtered dataset contains 768 loci with an average of 347 bp, 25 samples, and 50 SNPs per locus and an average of 107 loci per sample for the ingroup. Only two outgroup samples passed the filtering.

### Phylogenetic inference

Our final concatenated tree shows a mean UFBS likelihood of 97.16% across all nodes. QC, QD and QI show a mean of 0.27, 0.50 and 0.72 across all nodes, respectively. QF shows a mean of 0.39 across the tips. The coalescent tree receives a mean PP of 0.73 and a mean QS score of 0.25/0.55/0.72 across all nodes (supplementary Fig. S4). QF shows a mean of 0.38 across the tips of the coalescent tree.

The genus *Kalanchoe* was reconstructed as monophyletic, and all species except for *K. pareikiana* Desc. & Lavranos were retrieved within one of the four clades in both the concatenated and coalescent tree, referred to as clades A, B, C, and D (Fig. 2, Fig. 3, supplementary Fig. S3, Fig. S4). In the concatenated tree (Fig. 2), *K. pareikiana* formed a sister to clade (B, (C, D) while the species was part of clade B in the coalescent tree (supplementary Fig. S4).

**FIG. 2.**
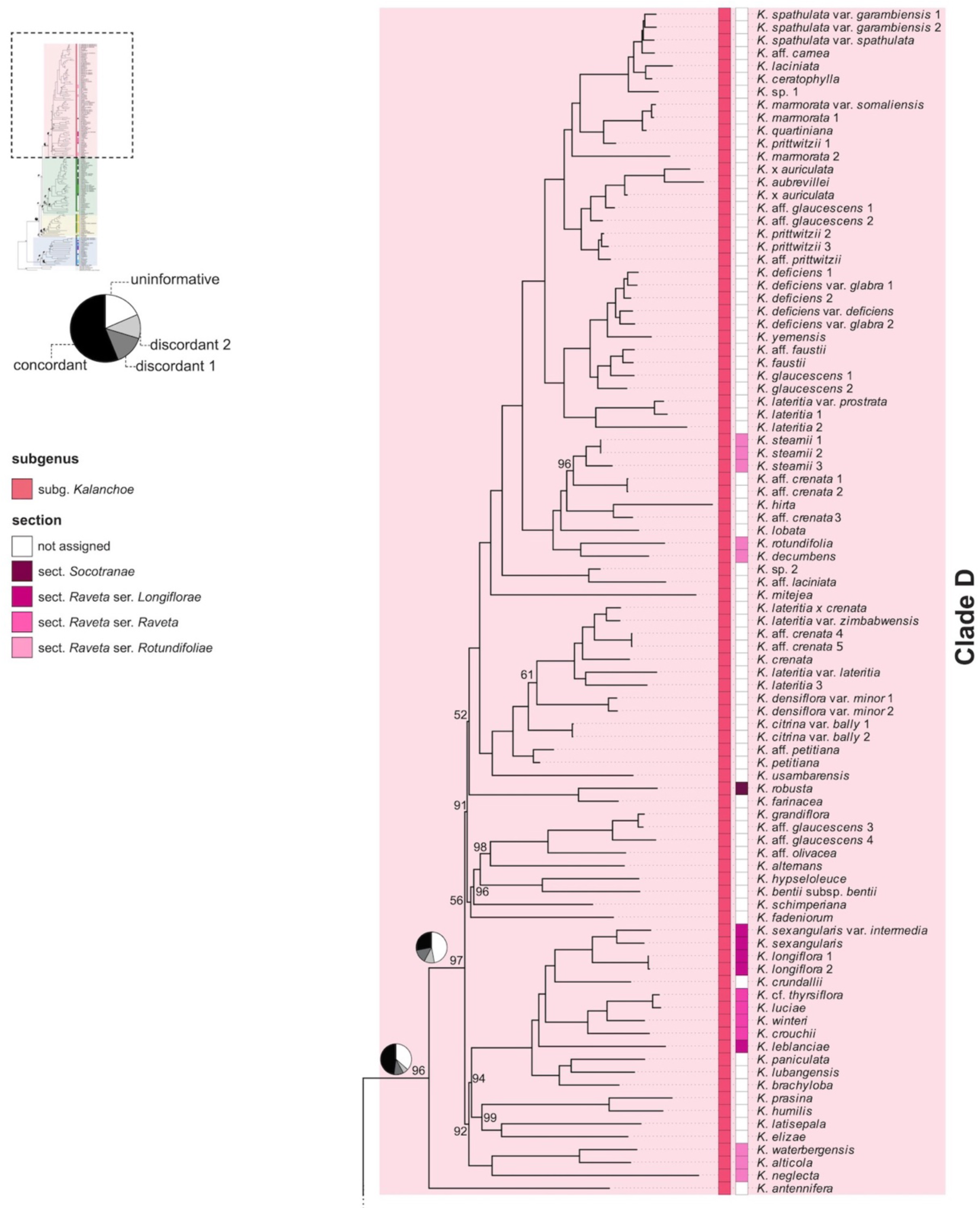
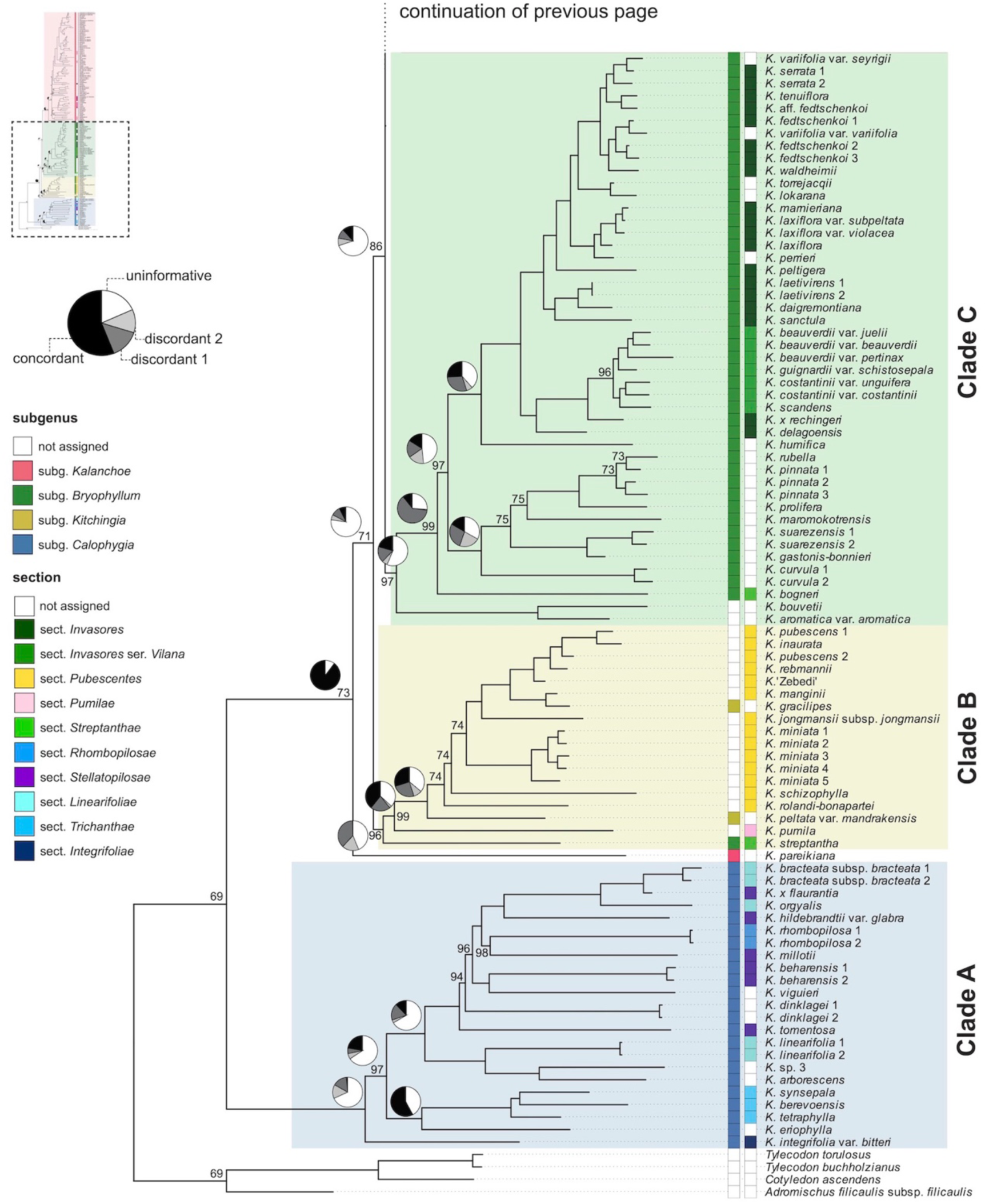
Maximum likelihood phylogeny of *Kalanchoe s.l.* inferred from the concatenated alignment of ddRADseq loci (concatenated tree). Colour highlighting of the tree corresponds to the four major clades. Coloured boxes represent the current classification system at the ranks of subgenus, section and series. Pie charts visualise the Quartet Sampling results for selected nodes in the backbone by showing the proportion of quartets that are concordant with the tree topology for this node (black), as well as the proportion of quartets supporting the two alternative, discordant topologies (light grey and dark grey) and the proportion of uninformative quartets (white). Bootstrap support values are shown only for nodes that did not receive maximum support. For full support values, see Fig. S4.

**FIG. 3.**
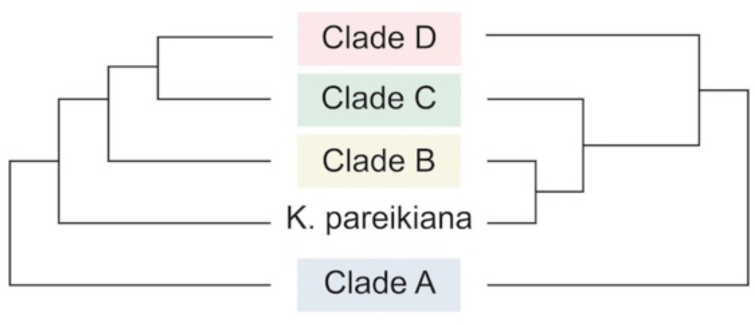
Schematic drawing of the backbones of the ddRADseq phylogenies of *Kalanchoe s.l.* inferred with a maximum likelihood approach from the concatenated alignment (concatenated tree; shown on the left) and with a two-step coalescent approach (coalescent tree; shown on the right).

In both analyses the first bifurcation separates Clade A from a sister clade comprising clade B, C and D and *K. pareikiana*. The monophyly of clade A, containing 23 samples representing 18 taxa, receives 100% UFBS support in the concatenated tree and a PP of 0.87 in the coalescent tree, while QS shows countersupport (concatenated tree: QC/QD/QI = −0.23/0.99/0.32, coalescent tree: −0.28/0.60/0.30) for this node. In the concatenated tree *K*. *integrifolia* Baker resolves as sister to the rest of clade A, which is again made up by two clades, one containing four species and the other 13 species, respectively. The alternative topologies, however, i.e. placing *K. integrifolia* as sister to the entire genus or as sister to the clade comprising clades B, C, and D and *K. pareikiana* receive stronger QS support. In the coalescent tree, *K. integrifolia* resolves in a polytomy with the same two clades that were recovered in the concatenated tree, containing four and 13 species respectively.

The monophyly of the combined clade B, C, and D including *K. pareikiana* is fully supported by QS (concatenated tree: 1/–/0.9, coalescent tree: 0.96/0.67/0.84), even though it receives relatively low UFBS support (73%) and PP (0.87) (Fig. 2, supplementary Fig. S3, Fig. S4). The relationship between clades B, C and D, however, differs between the analyses. In the concatenated tree, clade B is sister to clade (C, D) while in the coalescent tree, clade B and C form a sister relation, as sister to clade D. (Fig. 3). In the concatenated tree, *K. pareikiana* resolves as sister to clade (B, (C, D)). The position of *K. pareikiana* as sister to clade (B, (C, D)) is, however, very uncertain. The monophyly of the clade (B, (C, D)) excluding *K. pareikiana* receives low UFBS support (71%) and a QS score of −0.03/0.71/0.23 (supplementary Fig. S3). This low QS score for the monophyly of clade (B, (C, D)), together with the full support for the clade when *K. pareikiana* is included, indicates that this species might be better placed within clade B, as it is recovered in the coalescent tree, or within one of the other clades C or D.

The sister group relationship of clade C and D, as it is resolved in the concatenated tree, is only weakly supported (UFBS = 86%; QS = 0.01/0.88/0.3). The alternative topologies of a clustering of clades (B, D) or clades (B, C) receive comparable support from QS (Fig. 2, supplementary Fig. S3), but the high number of uninformative quartets indicates a high uncertainty for this node. In the coalescent tree, clades B and C resolve in a sister group relationship but with similarly high uncertainty. PP shows very weak support for the clustering of clades B and C (0.31) and QS shows countersupport (−0.11/0.84/0.26) but again based on a low number of informative quartets.

The monophyly of clade B, containing 18 samples representing 13 species, receives 96% UFBS support and a QS score of −0.38/0.65/0.56 in the concatenated tree. QS thus indicates that the placement of *K. streptantha* Baker, which resolves as sister to clade B, is highly uncertain. In the coalescent tree, the backbone within clade B shows a polytomy of 3 lineages: the first containing *K. pareikiana*, the second *K. pumila* and *K. peltata* and the third containing the remaining 10 species.

The monophyly of clade C, containing 46 samples representing 37 taxa, receives moderate support (concatenated tree: UFBS = 97; QS = 0.1/0.51/0.44, coalescent tree: PP = 0.78, QS = 0.15/0.44/0.58). Within clade C, a small clade containing *K. aromatica* and *K. bouvetii* resolves as sister to the rest of clade C in both analyses. Furthermore, in the concatenated tree, *K. bogneri* Rauh resolves as sister to two clades containing 27 and seven taxa respectively with 99% UFBS support. QS, however strongly favours the clustering of *K. bogneri* with *K. aromatica* and *K. bouvetii*. In the coalescent tree, the same two clades are recovered except that *K. bogneri* resolves as sister to the second clade in the coalescent tree, and *K. humifica* Desc. is recovered in a polytomy with the two clades, while it resolves as sister to the rest of the first clade in the concatenated tree.

The monophyly of clade D, containing 92 samples representing 70 taxa is reasonably well supported (concatenated tree: UFBS = 96, QS = 0.35/0.72/0.64, coalescent tree: PP = 0.99, QS = 0.15/0.76/0.32). Within clade D, branch lengths in the backbone are short, and the relationships among clades are uncertain in both analyses.

## Discussion

### Resolving the phylogeny of Kalanchoe

We generated the most densely sampled phylogeny of *Kalanchoe* to date, covering 70% of all taxa in the genus. The phylogenetic framework sheds new light on the relationships of the major groups within the genus, and their potential evolutionary processes. We identify four major clades in both our concatenated and coalescent analyses, herein referred to as clades A, B, C, and D (Fig. 2, Fig. 3, supplementary Fig. S3, Fig. S4). Both trees recover clade A as sister to the rest of *Kalanchoe*, while the relationships among the other clades are less certain. The concatenated tree resolves a weakly supported sister relationship of clades C and D, with QS showing comparable support for the two alternative topologies of a sister relationship of clades (B, D) or (B, C). In the coalescent tree, clades B and C are recovered as sister to each other. As the concatenated analysis includes all available information, the inferred phylogeny can be regarded as relatively robust. The coalescent analysis, on the other hand, is influenced by limited information contained in individual ddRAD loci. We thus herein focus on the results of the concatenated analysis even though the higher number of shared loci between individual samples of clades B and C, as compared to the number of shared loci between samples of either of the clades B or C with clade D (supplementary Fig. S2), additionally supports the results of the coalescent analysis, placing clades B and C as sister to each other.

The uncertainty in the relationships of clade B, C, and D could be influenced by the low number of loci shared across the backbone of the tree and, furthermore, these loci being shared by a comparatively low number of samples. Throughout evolutionary time, ddRAD loci dropout can occur through either the mutation and loss of cut sites or the gain of new cut sites, thus disrupting ancestral loci, while new loci that are shared among descendant clades arise (Eaton *et al*., 2016). Accordingly, our dataset shows a decreasing number of loci shared between individual samples with increasing phylogenetic distance (supplementary Fig. S2). For example, only two samples within the earliest diverging clade A, (*K. beharensis* 2 and *K. viguieri*) share more than ten loci with any sample from the other clades. We suggest that this contributes to the low QS scores across deeper nodes in our trees. However, we argue that quartet sampling can still provide valuable information on the stability of inferred relationships when carefully considered. While the number of shared loci between individual samples decreases with phylogenetic distance, the total number of shared loci across a branch is influenced by tree structure. Balanced trees with species-rich lineages, such as ours, are buffered against loci dropout through hierarchical redundancy, as mutations occurring more recently in evolutionary time only affect the descendants of the lineage where the mutation occurred, while the locus is still shared among other lineages (Eaton *et al*., 2016). As such, while each locus is only present in a reduced number of samples, the total number of loci recovered for a given branch can still be sufficient for robust phylogenetic inference. For example, across the deepest split within the genus *Kalanchoe* in our phylogeny, i.e., between clade A and the rest of the genus, 286 loci are shared, which corresponds to approximately 1% of all loci in the dataset. Therefore, we suggest that our tree still harbours sufficient data to infer phylogenetic relationships, and that although QS values may be affected by lower amounts of data at some nodes, they are still a meaningful, albeit general, reflection of discordance.

We suggest that some of the discordance in the relationship among clades B, C, and D could be a real biological signal resulting from hybridisation or introgression among species or ancestral lineages. Hybridisation events, both recent and ancient, are, in fact, highly probable to have occurred, at least among clades B and C. Species from clades B, C, and D, are capable of hybridisation across clades, but pre- and post-fertilisation incompatibilities apparently exist with species from clade A (RS and GFS pers. obs.). While clades B and C are indigenous to Madagascar, clade D is largely geographically isolated from the other clades, being primarily distributed in mainland Africa, Arabia, and Asia with few species occurring on Madagascar. As such, although they are sexually compatible, there has probably been little opportunity for hybridisation between species of clade D and other clades in the wild. Future work with target-capture or transcriptomic data could test our hypothesis of hybridisation events between clades B and C.

### Taxonomic implications

Clades A, B, C, and D are broadly congruent with the current subgeneric classification of *Kalanchoe*, with some exceptions. Clades A and D correspond to *K.* subg. *Calophygia* and *K.* subg. *Kalanchoe*, respectively, and are treated first in the discussion that follows. Clades C and B contain *K.* subg. *Bryophyllum* and *K.* subg. *Kitchingia*, respectively, as well as species that currently lack a subgeneric assignment.

#### Clade A: Kalanchoe subg. Calophygia

Clade A corresponds to the current circumscription of *K.* subg. *Calophygia*, also known as the ‘woody clade’ (Fig. 1 A–C). This subgenus was recently reinstated by Smith (2023*f*), and its circumscription was amended substantially from the original definition of the subgenus *sensu* Descoings (2006), who had additionally included species from all other clades in it (supplementary Fig. S5). Currently, the subgenus comprises 22 species and nothospecies, 17 of which were sampled for this study. The monophyly of *K.* subg. *Calophygia sensu* Smith (2023*f*) and its phylogenetic placement as sister to the remainder of *Kalanchoe* is consistent with the findings of Han *et al*., (2024) based on plastome sequence data. In the ITS-based phylogeny of Gehrig *et al*. (2001), what was later described as *K.* subg. *Calophygia* also resolved as monophyletic, although not as sister to the rest of the genus but as sister to a clade made up of species from *K.* subg. *Kalanchoe*. This corresponded to the taxonomic classification used throughout most of the taxonomic history of the genus, where these two clades were grouped together in the autonymic taxon *Kalanchoe* (treated at various ranks). The low resolution in the backbone of the tree of Gehrig *et al*. (2001), however, indicates low confidence in their topology and our results clearly show that these two lineages are not closely related within *Kalanchoe*.

Morphologically, *K.* subg. *Calophygia* and *K.* subg. *Kalanchoe* show similarities in flower morphology. Representatives of both subgenera have multidirectional or erect flowers and often (almost) free sepals, have the filaments inserted in the upper half of the corolla tube and the style shorter than the ovaries. *Kalanchoe s*ubg. *Calophygia* is, however, distinguished from the other subgenera by containing woody species with a pseudo-rosulate, shrubby, or arborescent habit, while the other subgenera do not contain truly woody species (Smith 2023*f*). Species from *K.* subg. *Calophygia* occur primarily in the dry southwest of Madagascar. Within the subgenus, five sections have recently been published (Smith 2020; 2023*b*; *e*; 2024*b*). These sections, with the exception of *K.* sect. *Rhombopilosae* Gideon F.Sm., are based on unranked groups proposed by Berger (1930). *Kalanchoe integrifolia*, included in the monotypic *K.* sect. *Integrifoliae* (A.Berger) Gideon F.Sm. (Smith, 2024*b*), resolves as sister to the rest of *K.* subg. *Calophygia* in the concatenated tree, although with some uncertainty in its placement (Fig. 2). Berger (1930) had characterised his *K.* [infragen.unranked] *Integrifoliae*, on which the current *K.* sect. *Integrifoliae* is based, by the absence of stellate hairs, entire leaf margins, and by the filaments being inserted in the lower half of the corolla tube (while “in the upper half of the corolla tube” would be more accurate, SB pers. obs.). *Kalanchoe* sect. *Trichanthae* (A.Berger) Gideon F.Sm., containing three species, resolves as monophyletic, clustering with *K. eriophylla* Hils. & Bojer ex Tul.. The three species included in *K.* sect. *Trichanthae* are characterised as low-growing, pseudo-rosulate perennials that are either stemless or with a short, sturdy, woody stem (Smith, 2023*e*). *Kalanchoe eriophylla* had been included in *K.* [infragen. unranked] *Stellatopilosae* by Berger (1930) based on the presence of stellate hairs but later has been excluded from *K.* sect. *Stellatopilosae* (A.Berger) Gideon F.Sm. by Smith (2023*f*). The third clade within *K.* subg. *Calophygia* is recovered as sister to *K.* sect. *Trichanthae* and *K. eriophylla* in the concatenated tree and contains what is today accepted under *K.* sect. *Stellatopilosae*, *K.* sect. *Linearifoliae* (A.Berger) Gideon F.Sm., and *K.* sect. *Rhombopilosae*. While *K.* sect.

*Rhombopilosae* is monotypic, the other two sections both resolve as polyphyletic. The main character used by Berger (1930) to distinguish his two unranked infrageneric groups, i.e., *K.* [infragen. unranked] *Linearifoliae* and *K.* [infragen. unranked] *Stellatopilosae*, on which the current sections are based, is the presence or absence of stellate hairs, which seems to be a variable character insufficient for sectional differentiation. Earlier, Boiteau and Allorge-Boiteau (1995) had proposed an informal infrageneric classification system for the clade. While their informal groups XII Trichanthae and XIII Integrifoliae are identical to the currently accepted *K.* sect. *Trichanthae* and *K.* sect. *Integrifoliae*, they grouped all of the other species in the informal group X Lanigerae, which resolves as monophyletic in both of our trees if *K. eriophylla* were excluded (see supplementary Fig. S5). While further research is needed to establish the taxonomic status of the sections currently recognised in *K.* subg. *Calophygia*, our tree gives strong support for the recognition of clade A as a natural subgenus of *Kalanchoe*.

#### Clade D: Kalanchoe subg. Kalanchoe

Clade D corresponds to the autonymic *K.* subg. *Kalanchoe* (Fig. 1 G, I). This finding is in contrast to previous molecular work, where in the ITS-based phylogeny of Gehrig *et al*. (2001) the species from this subgenus resolved as polyphyletic. However, the low support of the backbone, renders their phylogenetic inference unreliable. Morphologically, the clade is distinguishable by having erect to multidirectional flowers, as *K.* subg. *Calophygia*, but with a narrow opening of the often long corolla tube. *Kalanchoe* subg. *Kalanchoe* is the only subgenus of *Kalanchoe* that is not endemic to Madagascar but it seems to have a Malagasy origin. *Kalanchoe antennifera* Desc., the only Malagasy representative of the subgenus sampled for this study, resolves as sister to the rest of the clade in the concatenated tree. *Kalanchoe* subg. *Kalanchoe* has a wide geographical distribution, naturally occurring mainly in southern and eastern continental Africa, and additionally on the Arabian Peninsula, and in Southeast Asia and northwest Australia. We thus hypothesise that the dispersal out of Madagascar only happened once, with subsequent diversification in continental Africa and dispersal to the Arabian Peninsula, Southeast Asia, and northwest Australia. While we did not conduct a formal biogeographic analysis, we nevertheless observe that the Southeast Asian species do not form a monophyletic clade but resolve in two separate clades within the subgenus, both in the concatenated and in the coalescent tree (Fig. 2, supplementary Fig. S4). The two Indian species, *K. grandiflora* Wight & Arn., and *K.* aff. *olivacea* Dalzell, resolve in a clade that otherwise contains species from northern Africa and the Arabian Peninsula, while the other Southeast Asian species, including the type of the genus, *K. laciniata* (L.) DC., resolve in a different clade, sister to species from mainly northern and eastern Africa. Similarly, species indigenous to the Arabian Peninsula were recovered in two distinct clades, one containing *K. alternans* (Vahl) Pers. and *K. bentii* C.H.Wright ex Hook.f. and the other *K. deficiens* (Forssk.) Asch. & Schweinf. and *K. yemensis* (Deflers) Schweinf., but both times clustering with species from northern Africa. Although *K.* subg. *Kalanchoe* is very species-rich, only a few infra-subgeneric classifications have been proposed. *Kalanchoe* sect. *Raveta* Raym.-Hamet ex Gideon F.Sm., currently contains 20 species indigenous to southern and south-tropical Africa (Smith, 2024*c*). The section is furthermore split into three series. The four species sampled from *K.* [sect. *Raveta*] ser. *Raveta* Gideon F.Sm. form a monophyletic clade within a larger clade of South African species, which additionally includes *K.* [sect. *Raveta*] ser. *Longiflorae* Gideon F.Sm. and three other species. *Kalanchoe* [sect. *Raveta* ser. *Rotundifoliae*] Gideon F.Sm. resolves as polyphyletic and separate from the rest of *K.* sect. *Raveta. Kalanchoe* sect. *Socotranae* (A.Berger) Gideon F.Sm. is a monotypic section indigenous to Socotra (Berger, 1930; Smith, 2024*b*).

Within *K.* subg. *Kalanchoe*, the relationships among the clades are not well resolved in our phylogeny and differ between the methods. This might be due to rapid diversification, potentially linked to dispersal across the continent, leading to short branch lengths and little information contained in each branch and, thus limiting the capability of the data to resolve relationships fully. On the species level, the phylogenetic analysis indicates the need for further taxonomic work, as several species resolve as polyphyletic. The clade D as a whole, however, forms a reliable natural unit, supporting its treatment at the rank of subgenus.

#### Clade C: Kalanchoe subg. Bryophyllum

Clade C contains two subclades, one consisting of *K. aromatica* and *K. bouvetii* and its sister clade corresponding to species that are currently treated in *K.* subg. *Bryophyllum*, including the type of the subgenus, *K. pinnata* (Lam.) Pers.. *Kalanchoe aromatica* and *K. bouvetii* currently lack a subgeneric assignment but *K. aromatica* has in the past mainly been classified under the taxon *Kalanchoe* (e.g. Berger, 1930; Boiteau and Allorge-Boiteau, 1995) while *K. bouvetii* has been treated under both *Bryophyllum* (Berger, 1930; Descoings, 2003) and *Kalanchoe s.s.* (Boiteau and Allorge-Boiteau, 1995). Both species had been included in *K.* subg. *Calophygia* by Descoings (2006) but were later excluded from it by Smith (2023*f*), but in contrast to most other species, they were not reassigned to another subgenus (Smith, 2024*a*). The species of the second subclade of clade C, sister to *K. aromatica* and *K. bouvetii*, have consistently been classified in the taxon *Bryophyllum* across most classification systems, treated at genus, subgenus, or section level (supplementary Fig. S5). In the ITS-based phylogeny of Gehrig *et al*. (2001), species from clade C resolved as monophyletic. Equally, the four species from *K.* subg. *Bryophyllum* included in the study by Han *et al*. (2024) formed a clade. Neither of the two studies, however, had included *K. aromatica* or *K. bouvetii*.

*Kalanchoe aromatica* and *K. bouvetii* differ from the rest of clade C in having multidirectional flowers with distinctly recurved corolla lobes, while the rest of the clade has pendulous flowers with the corolla lobes sometimes pointing outward but not being recurved. Furthermore, *K. aromatica* and *K. bouvetii* lack the ability to produce bulbils on the leaf margin. This trait is present in species of the sister clade with the exception of *K. bogneri* (Fig. 1 D) which branches off first in this clade, at least in the tree based on the concatenated matrix (Fig. 2). This indicates that the phyllo-bulbiliferous species evolved only once in the genus and are restricted to this clade, while bulbils in the inflorescence and other vegetative reproductive strategies are more widespread and scattered across the genus (Smith *et al*., 2022). Within this leaf-bulbil-producing clade, *K.* sect. *Invasores* Shtein & Gideon F.Sm. resolves as a well-supported and monophyletic clade both in the concatenated and the coalescent tree. Within *K.* sect. *Invasores*, the climbing species, i.e. *K.* ser. *Vilana* Shtein & Gideon F.Sm., resolves as monophyletic. *Kalanchoe* sect. *Invasores*, together with *K. humifica* in the concatenated tree, is sister to a clade that corresponds to the informal group IX Proliferae as defined by Boiteau and Allorge-Boiteau (1995), which was, however, not validly published. In the coalescent tree *K. humifica* is recovered in a polytomy with the two other clades.

While all species of clade C (both subclades) are indigenous to Madagascar, some members, especially from *K.* sect. *Invasores*, have been introduced to other parts of the world. Probably enhanced by their ability to reproduce vegetatively through bulbils, some species have become invasive in areas with a Mediterranean, subtropical, and tropical climate (Herrando-Moraira *et al*., 2020; Shtein and Smith, 2021; Smith, 2023*d*).

#### Clade B: Kalanchoe subg. Kitchingia and K. sect. Pubescentes

Clade B, as recovered in the concatenated tree, includes at subgenus rank *K.* subg. *Kitchingia* and one species currently treated in *K.* subg. *Bryophyllum.* Most other species from clade B, however, match the description of the recently published *K.* sect. *Pubescentes* (Smith, 2024*d*; GFS unpublished data), which has not been assigned to a subgenus. *Kalanchoe pareikiana*, which falls within clade B in the coalescent tree but resolves as sister to the combined clade (B, (C, D)) in the concatenated tree, is currently treated in *K.* subg. *Kalanchoe*.

No previous taxonomic classification system has treated all of clade B as a unit, but earlier treatments have variously grouped subsets of species contained in the clade, treating them under *Kitchingia*, *Bryophyllum*, or *Kalanchoe* at various ranks. Morphologically, clade B is diverse and rather challenging to differentiate from the other clades, which is also reflected in the complex taxonomic history of the clade (see Fig. 2, supplementary Fig. S5). The flowers are mostly pendulous, with the exception of *K. pumila* and *K. jongmansii* Raym.-Hamet & H.Perrier, a trait shared with most species of clade C. Furthermore, most species, except for *K. streptantha*, have a short calyx with sepal lobes longer than the calyx tubes. With the exception of *K. pumila*, the species have short ovaries combined with a long style. Many species in the clade form dense clusters of bulbils in the post-anthesis inflorescence. This trait is, however, not restricted to clade B. Species from clade B are endemic to Madagascar and occur mainly in the central highlands as well as in the more humid northern and eastern parts of the island. Our tree indicates the need for taxonomic work on the classification of clade B in order to render subgenera monophyletic. The tree supports the treatment of clade B as a natural unit.

In the ITS-based phylogeny of Gehrig *et al*. (2001), species from our clade B were, with the exception of *K. streptantha*, recovered in two distinct clades. They treated the first clade as section *Kitchingia*, containing the two species currently accepted in *K.* subg. *Kitchingia*—*K. gracilipes* and *K. peltata.* The second clade was included in their *K.* sect. *Bryophyllum* and contained all other species they had sampled from clade B, except for *K. streptantha*, which was recovered in a clade with species from clade C in their tree. In the study by Han *et al*. (2024), none of the species from clade B were included.

The current definition of *K.* subg. *Kitchingia sensu* Smith *et al*. (2021*a*) comprises only the two species *K. gracilipes* and *K. peltata*. Both species have fully divergent carpels and differ from other campanulate-flowered species by lacking dark vein patterns on the inside of the corolla (Smith *et al*., 2021*a*). Throughout the taxonomic history of the genus, the two species have mostly been treated under the taxon *Kitchingia*, albeit at different ranks and with a different set of species included in the taxon. The subgenus resolves, however, as polyphyletic in both the concatenated and coalescent trees. A wider definition of the subgenus *Kitchingia*, which is still resolving as polyphyletic in both our trees, was used by Smith and Figueiredo (2018). They additionally included *K. miniata* Hils. & Bojer ex Tul.*, K. schizophylla* Baill. & H.Perrier, *K. ambolensis* Humbert*, K. campanulata* Baill., *K. porphyrocalyx,* and *K. uniflora* in the subgenus. The last four of these species were not sampled in this study. *Kalanchoe miniata*, *K. schizophylla*, *K. ambolensis,* and *K. campanulata* currently belong to *K.* sect. *Pubescentes* (GFS unpublished data) and lack subgeneric classification while *K. porphyrocalyx* and *K. uniflora* are treated in their own subgenus, *K.* subg. *Alatae* (Smith, 2023*a*).

The remaining species within our clade B, with the exception of *K. pumila*, and *K. streptantha*, are grouped together in the recently described *K.* sect. *Pubescentes* (GFS unpublished data). Representatives of this section are mostly characterised by often being at least sparsely pubescent, creeping to leaning to erect, succulent subshrubs often with hard-wiry to flexuose to rigid stems; by their bulbiliferous inflorescences with the bulbil clusters that are dense and long-lasting; and by the sepals that are fused for ± ⅓ to ½ of their lengths. The section resolves as polyphyletic with *K.* subg. *Kitchingia* falling within the clade.

*Kalanchoe streptantha*, which resolves as sister to the rest of Clade B in our concatenated tree, is currently treated in *K.* [subg. *Bryophyllum*] sect. *Streptanthae* Gideon F.Sm., containing *K. streptantha* and *K. bogneri*, a species from clade C (Smith, 2023*c*). The section thus resolves as polyphyletic. The characters used by Smith (2023*c*) to define the section include the plants being non-bulbiliferous, the calyx being substantially fused, the cylindrical corolla being constricted toward the middle (traits shared with other species of clades B and/or C), and the pedicels enlarging toward the calyx (which can however not be observed in *K. bogneri* in this form). Our tree does not support the grouping of the two species in a section. *Kalanchoe pumila* (Fig. 1 F), together with *K. bergeri* Raym.-Hamet & H.Perrier which was not sampled for this study, is currently treated in *K.* sect. *Pumilae* (A.Berger) Gideon F.Sm., which was published recently based on an unranked group of Berger (1930; see Smith, 2024*b*). The species included in this section currently have no subgenus assigned. The third species treated in *K.* [infragen. unranked] *Pumilae* by Berger (1930), *K. jongmansii*, is currently interpreted as belonging to *K*. sect. *Pubescentes* (GFS unpublished data).

### Implications for the evolution of Kalanchoe

Our phylogenetic tree allows new hypotheses about the evolution of *Kalanchoe* and provides a framework for future research to test them. The production of bulbils on the leaf margin (Fig. 1 E) is restricted to a large clade within *K.* subg. *Bryophyllum*, which constitutes most of clade C. The production of bulbils in the inflorescence (Fig. 1 K), on the other hand, is more widespread in the genus, even though it is most pronounced in species from clades B and C. We might thus hypothesise that the production of bulbils on the leaf margin could be based on a more specific and complex genetic mechanism, while vegetative reproduction from other parts of the plants, especially from meristems already present in the inflorescence following its branching pattern, might be easier to achieve and is in fact also shared with other representatives of Crassulaceae (Berger, 1930). Jácome-Blásquez and Kim (2023) conclude from transcriptomic and immunolocalization studies that for bulbil formation on the leaf margin in *Kalanchoe* key genes involved in meristem formation and development are co-opted in a new context to acquire meristem competency.

Our tree could also have interesting implications for the evolution of CAM photosynthesis. It had been hypothesised based on stable isotope values that some of the strongest levels of CAM were expressed by species now included in *K.* subg. *Calophygia* while species from *K.* subg. *Kitchingia* were described as displaying almost C_3_-like values (Kluge *et al*., 1991; 1993). A genus-wide study of CAM expression based on thorough physiological measurements is, however, lacking to date, despite some species having been studied in depth and used as model species for CAM photosynthesis (e.g. Hartwell *et al*., 2016; Boxall *et al*., 2017; Yang *et al*., 2017; Winter, 2019). The phylogenetic tree can thus provide a framework to expand our knowledge on CAM photosynthesis by studying its evolution in depth across the genus. Messerschmid *et al*. (2021) compiled 112 carbon isotope ratio measurements for *Kalanchoe* (mainly from Kluge *et al*., 1991; 1993) ranging from −30.6 to −10.2 indicating a strong diversity in CAM expression in the genus, even within several species (e.g. *K. porphyrocalyx* −30,6 to - 15,7 (N=7) and *K. lanceolata* (Forssk.) Pers. −23,6 to −12,0 (N=10)). The majority of the 56 species investigated so far seem to express strong CAM, however often only one sample was measured.

Our phylogeny also sheds new light on the biogeographic history of the genus. *Kalanchoe* subg. *Kalanchoe* resolves as monophyletic, and it is the only subgenus not endemic to Madagascar. This suggests a Malagasy origin of the genus as a whole, as was hypothesised by Allorge-Boiteau (1996). Although we only sampled one Malagasy representative of *K.* subg. *Kalanchoe*, we hypothesise that the dispersal to the African mainland only happened once, with subsequent dispersal on the continent. The biogeographic patterns on the African mainland itself, as well as the dispersal to the Arabian Peninsula and Southeast Asia, however, require a more thorough investigation.

Our phylogenetic tree can further be used to study the evolution of morphological traits, especially flower morphology, leaf morphology, and growth form. Clade B and clade C seem to be morphologically more similar to each other than to clade A or D, as reflected in the taxonomic history of the genus, but they do not form a clade together in our concatenated tree. Some uncertainty remains, however, in the relationships between clades B, C, and D, suggesting the need for further phylogenetic studies using complementary sequencing technologies, as for example phylotranscriptomics or target capture, that allow for the investigation of (ancient) hybridisation and introgression among the major clades, as this could have an influence on tree inference. The four major clades themselves nevertheless form stable units.

### Conclusion

This study has advanced our understanding of the relationships within the genus *Kalanchoe*. Our phylogenetic tree supports the treatment of the *K.* subg. *Calophygia* and *K.* subg. *Kalanchoe* as currently recognised but suggests that *K.* subg. *Bryophyllum* and *K.* subg. *Kitchingia* are not monophyletic in their current circumscription and warrant taxonomic revision. Future studies should focus on further improving the data by the use of complementary sequencing methods that additionally allow the investigation of deep reticulation among clades. Future studies should also include *K.* subg. *Alatae*, which is missing from our taxon sampling. Nevertheless, as it stands, our phylogeny raises interesting hypotheses about the evolution of traits, such as CAM photosynthesis, flower and leaf morphology, growth form, habitat requirements, and reproductive strategies, and provides a framework for their investigation.

## Data availability statement

Sequence data will be uploaded upon acceptance of the paper

## Acknowledgements

Silvia Wienken and Alina Höwener, Agnes Scheunert, and Om Kulkarni are thanked for their support and advice with lab work and sequence alignment. The Botanischer Garten München Nymphenburg is thanked for hosting and curating a comprehensive living collection of *Kalanchoe*, in particular Franziska Berger, Betina Seisenberger, Franziska Schustetter, David Ullmann, Sebastian Walter (all BG Munich) and Christopher Wild (BG Mainz) are thanked for taking care of the plants. A substantial part of the Munich collection came from the Botanical Garden Mainz in 2021. The following botanic gardens are thanked for sharing cuttings from their living collection: The Yehuda Naftali Botanic Garden of Tel Aviv University (TELA), Botanischer Garten der Universität Heidelberg (HEID), Sukkulenten-Sammlung Zürich, Palmengarten der Stadt Frankfurt (FRP), Jardin Botanique de la Mairie de Lyon (LYJB), Botanischer Garten der Universität Düsseldorf (DUSS). Botanischer Garten und Botanisches Museum Berlin-Dahlem (B), and Muséum national d’Histoire naturelle in Paris (MNHN) are thanked for providing silica dried material from their living collection. Alessandra Havinga and Louis Nusbaumer are thanked for providing a silica dried sample collected in Madagascar. Richard Razakamalala and Faranirina Lantoarisoa from Missouri Botanical Garden as well as the herbarium TAN are thanked for assistance and collaboration during fieldwork in Madagascar. Leonie Hinderhofer is thanked for help with graphical design of Figures 2 and 3. The Ludwig-Maximilians-Universität München is thanked for funding the project as part of the seed funding provided to GK.

## Author contributions

SR designed the study, conducted fieldwork and lab work, established lab protocols, analysed the data and wrote the manuscript. DPK, RS and GFS provided and identified samples, conducted fieldwork, provided insight through fruitful discussions. EJ and DMB provided advice on data analysis and writing of the manuscript. HM conducted lab work, in particular establishing lab protocols for *Kalanchoe*. RL conducted fieldwork. PH established lab protocols and designed adapters. SB provided and identified samples. GK designed the study, acquired funding and provided advice on writing of the manuscript. All co-authors revised the manuscript.

## SUPPLEMENTARY MATERIAL

Supplementary Table S1. Accession table of specimens sampled.

Supplementary Figure S1a. Comparison of within-sample clustering thresholds for four metrics that were evaluated for the assembly with the Ipyrad pipeline. The total number of clusters, total average depth, loci filtered by maximum heterozygosity, and heterozygosity per sample are shown across within-sample clustering thresholds of 80–98% similarity between reads.

Supplementary Figure S1b. Comparison of between-sample clustering thresholds for four metrics that were evaluated for the assembly with the Ipyrad pipeline. The total number of retained loci, the total number of variable sites, the percentage of missing data and the average number of variable sites per locus are compared across between-sample clustering thresholds of 80–98% similarity between within-sample consensus sequence clusters.

Supplementary Figure S2. Number of ddRADseq loci shared between samples. Rows and columns represent individual samples. They are ordered according to their placement in the concatenated tree, shown on the left. The diagonal entries show the number of loci recovered for each sample and off-diagonal entries represent the number of shared loci between the samples. The colour coding is log-scaled with darker colours representing more shared loci between samples.

Supplementary Figure S3. Quartet sampling scores on maximum likelihood phylogeny of *Kalanchoe s.l.* inferred from the concatenated alignment of ddRADseq loci (concatenated tree). Quartet sampling scores (Quartet concordance (QC), quartet discordance (QD) and quartet informativeness (QI)) are shown for each branch separated by a vertical line.

Supplementary Figure S4a. Quartet sampling scores on ddRADseq phylogeny of *Kalanchoe s.l.* inferred with a two-step coalescent approach (coalescent tree). Quartet sampling scores (Quartet concordance (QC), quartet discordance (QD) and quartet informativeness (QI)) are shown for each branch separated by a vertical line.

Supplementary Figure S4b. Local posterior probability support values on ddRADseq phylogeny of *Kalanchoe s.l.* inferred with a two-step coalescent approach (coalescent tree). local posterior probability support values are shown only for nodes that do not receive maximum support

Supplementary Figure S5. Partial taxonomic history of *Kalanchoe s.l.*: Comparison of selected taxonomic classifications from 1930 to the present in relation to the concatenated tree. Berger (1930) treated *Kalanchoe s.s.*, *Bryophyllum*, and *Kitchingia* as separate genera and divided *Kalanchoe* into 10 unranked units largely based on the numbered groups of Hamet (1907; 1908). Boiteau and Allorge-Boiteau (1995) recognised three sections in *Kalanchoe s.l.*: *Kitchingia*, *Bryophyllum*, and *Kalanchoe*. They furthermore split the genus into 15 informal groups. Descoings (2003) recognised two sections only in *Kalanchoe s.l.*: *Kalanchoe* and *Bryophyllum*. Affinities of species intermediate between the two sections were so indicated. Descoings (2006) recognised three subgenera in *Kalanchoe*: *Kalanchoe*, *Bryophyllum*, and *Calophygia*, with those species that he considered to be intermediate between *K.* subg. *Kalanchoe* and *K.* subg. *Bryophyllum* contained in *K.* subg. *Calophygia*. Smith and Figueiredo (2018) published a combination for *Kitchingia* at the rank of subgenus and Smith (2021) published the name *K.* subg. *Fernandesiae*. The circumscription of these two taxa has since been superseded. The last two columns represent the most recent taxonomic treatment at the ranks of subgenus and section (including series), largely representing the work of one of us (GFS) and colleagues since 2018. Smith (2023*g*; 2024*a*) provide comprehensive reviews of the infrageneric taxonomic history of *Kalanchoe s.l*.

