## Supplementary_Material for "A new phylogenetic framework for the genus *Kalanchoe* (Crassulaceae) and implications for infrageneric classification"

### Supplementary Table S1. Accession table of specimens sampled.

| Tip label | accession_number | Herbarium_voucher | collector | collection_date | Country | locality | comment |
|---|---|---|---|---|---|---|---|
| Adromischus filicaulis subsp. filicaulis | Berlin 278388320 | tba | NA | 29/10/1983 | Namibia | Lüderitz (District), 12 km N Rosh Pinah, steiniger Hügel mit Quarzbändern, ca, 600m. | NA |
| Cotyledon ascendens | Zürich 89 2809 /1 E. van Jaarsveld 9915 | tba | E. van Jaarsveld | 14/04/1989 | SouthAfrica | Cape Prov., Port Elizabeth, (3325 DC) Swartkops River. From the locus classicus! | received from Jaarsveld 24.7.1989 |
| K. aff. carnea | München 2000/1464 | tba | NA | NA | NA | NA | NA |
| K. alternans | XX-0-TELA-2021.0029 RS 170 Lav 30736 | tba | NA | NA | Yemen | Wadi Salul, Mukeiras, S Yemen | NA |
| K. alticola | XX-0-B-1940311 | tba | NA | NA | NA | NA | NA |
| K. antennifera | Palmengarten Frankfurt 6-22818-1 RMM 450 | tba | Ralph Mangelsdorff | 1998 | Madagascar | Antsalova, Tsingy de Bemaraha | NA |
| K. arborescens | München 2021/1480 Heidelberg 143206 Rauh 68581a | tba | W. Rauh | 08/06/1987 | Madagascar | NA | Donor: W. Rauh, 1987-08-06 |
| K. aromatica var. aromatica | München 2021/1534 Zürich 10 1913/0 Sieder et al 4893-1 | tba | Sieder et al. | 2009 | Madagascar | Antananarivo | NA |
| K. aubrevillei | München 2021/0727 Lyon 210033 | tba | NA | NA | Tanzania | Sud de Oldeani; WP 812 | NA |
| K. x auriculata 1 | XX-0-B-1459874 Bally 13469 | tba | Bally | 05.1970 | Kenya | NA | NA |
| K. beauverdii var. beauverdii | XX-0-TELA-2020.0461 RS 391 | tba | P. Richaud | 1994 | Madagascar | Cap Sainte Marie | NA |
| K. beauverdii var. juelii | RS 921 ISI 2018-24 | tba | Vincent Cerutti | NA | NA | NA | ex Jardin exotique de Monaco |
| K. beauverdii var. pertinax | G.F. Smith 1126 | PRU 127929 | NA | 1996 | Madagascar | NA | Ex hort. South Africa. Gauteng Province, Pretoria. Sample obtained from material from which the holotype of the name was preserved |
| K. beharensis 2 | XX-0-TELA-2021.0032 RS 814 | tba | Richaud | 1994 | NA | NA | NA |
| K. beharensis 1 | München 2021/1481 Heidelberg 140508 Rauh M1783 | tba | W. Rauh | 1959 | Madagascar | NA | Donor: W. Rauh, 1960-00-00 |
| K. bentii subsp. bentii | XX-0-TELA-2021.0033 Lav30753 RS 174 | tba | NA | NA | Yemen | Mola Matar, Hadhramant | NA |
| K. berevoensis | München 2021/1707 PalmengartenFrankfurt 18-30314-4 | tba | Norbert Rebmann | NA | NA | Anosiampela | Typmaterial |
| K. bogneri | München 1990/0786 Bogner 2138 | tba | Bogner | NA | Madagascar | Tsingy de Bemaraha, TYPUS | NA |
| K. bouvetii | München 2021/1533 Heidelberg 142376 | tba | NA | NA | Madagascar | Toliara Province, NW of Mahaboboka, Analavelona, forested mountain-chain, small brook in very steep and deep gorge, growing under shady conditions, above a boulder in | NA |
| K. brachyloba | G.F. Smith1136 | PRU 128249 | G.F. Smith | 1999 | Angola | Namibe province, Huíla plateau | NA |
| K. bracteata subsp. bracteata 2 | München 1994/0366 | tba | NA | NA | Madagascar | NA | NA |
| K. bracteata subsp. bracteata 1 | München 2021/1536 Heidelberg 101374 W.Rauh 74338 | tba | W. Rauh | 31/03/1994 | Madagascar | Toliara Province, Esomony, Granitfelsen | Donor: W. Rauh, 1994-00-00 |
| K. miniata 5 | München 2021/1537 Zürich 99 9305/0 | tba | Kluge | 1988 | Madagascar | Antananarivo | ex BG Darmstadt |
| K. miniata 4 | München 2021/1552 Zürich 15 0278/0 | tba | Röösli & Hoffmann | 1995 | Madagascar | NA | NA |
| K. ceratophylla | XX-0-TELA-2021.0035 RS 676 | tba | NA | NA | Thailand | South Thailand | NA |
| K. citrina var. bally 2 | Zürich 2014/0940 | tba | NA | NA | Kenya | Longopito | NA |
| K. citrina var. bally 1 | München 2014/1237 | tba | NA | NA | Madagascar | NA | NA |
| K. aff. crenata 5 | München 2021/1539 Zürich 99 5775/0 Pfennig 1052 | tba | Pfennig | 1972 | Kenya | Central Prov. | NA |
| K. costantinii var. costantinii | XX-0-TELA-2020.0456 RS 901 | tba | Richaud | 1994 | Madagascar | Berenty | NA |
| K. costantinii var. unguifera | XX-0-TELA-2020.0460 RS 957 | tba | Richaud | NA | Madagascar | Tolagnaro | NA |
| K. aff. crenata 3 | G.F. Smith 1138 | PRU 128251 | G.F. Smith | 1984 | SouthAfrica | KwaZulu-Natal | NA |
| K. aff. crenata 2 | München 2021/1542 Berlin 146037920 | tba | NA | NA | Kenya | NA | NA |
| K. aff. crenata 4 | München 2021/1540 Zürich 99 5796/0 Raadts & Nuernbergk 39 | tba | Raadts & Nuernbergk | 1968 | Kenya | Rift Valley Prov. | NA |
| K. crouchii | G.F. Smith 1139 | PRU 128252 | G.F. Smith | 2001 | SouthAfrica | Mpumalanga province, Pilgrim's Rest | Sample obtained from material from which the holotype of the name was preserved |
| K. crundallii | ZA-0-B-1340111 | tba | NA | NA | SouthAfrica | Limpopo (Prov.), Hanglip, Soutpansberg | NA |
| K. curvula 2 | München 2021/0731 Lyon 210042 RS 620 | tba | Ikeda | NA | Madagascar | Atsimo-Atsinanana, Ankaizina | JB Tananarive 11 |
| K. curvula 1 | SER23-014 | tba | Seraina E. Rodewald, David-Paul I | 05/10/2023 | Madagascar | road RN31, way to Bealanana | |
| K. daigremontiana | RS 473 | tba | Richaud | 1994 | Madagascar | Makay massif | NA |
| K. decumbens | XX-0-B-1880307 | tba | NA | NA | NA | NA | NA |
| K. glaucescens 2 | München 2021/0712 Lyon 210045 DPK 661 | tba | NA | NA | Ethiopia | NA | NA |
| K. deficiens 2 | München 2021/0713 Lyon 210046 | tba | NA | NA | Yemen | NA | NA |
| K. deficiens var. glabra 2 | RS 201 ISI 92-52 HBG 65307 | tba | NA | NA | NA | NA | NA |
| K. deficiens var. glabra 1 | YE-0-B-2330286 Deil 635 | tba | Deil | 06/10/1982 | Yemen | Hajjah (Prov.), Dorfrand von Mabyan, Kalk, 1800m | NA |
| K. densiflora var. minor 2 | KE-0-B-0239674 | tba | Hanid, Raadts & Nuernbergk | NA | Kenya | NA | NA |
| K. densiflora var. minor 1 | KE-0-HEID-45136 Rauh Ke488 | tba | Rauh | 13/02/1969 | Kenya | Rift Valley Province, [Bonde la Ufa], District Nakuru, oberhalb Naivasha, LatLong1: -0.6646580, 36.6400340. | Typus |
| K. dinklagei 2 | München N/3117 | tba | NA | NA | NA | NA | NA |
| K. dinklagei 1 | München 2021/1485 Heidelberg 141009 | tba | M. Tessier | NA | Madagascar | Androy Region | NA |
| K. elizae | XX-0-B-2301074 | tba | NA | NA | NA | NA | NA |
| K. eriophylla | München N/0172 | tba | NA | NA | NA | NA | NA |
| K. fadeniorum | KE-0-B-0240474 Faden & Faden 77/777 | tba | Faden & Faden | 11/03/1977 | Kenya | Kwale District, Mombasa - Nairobi road, 4 km towards Mombasa from turn off to Mackinnon Road Railway Station, semi-evergreen thicket on brown sandy soil with superficial patches of clayey soil in lower-lying spot, ca. 360m (L: 39°0'E / B: 3°44'S) | Samm.- Notes: Typus-Aufsammlung! (vgl. Holotypus B100241966; Sammeldaten dem Protolog entnommen) |
| K. farinacea | München 2021/1544 Heidelberg 140826 | tba | NA | NA | Yemen | Socotra | Donor: Les Cèdres, 1990-06-10 |
| K. faustii | XX-0-TELA-2021.0038 JAA202 RS 886 | tba | NA | NA | Morocco | Qued Quarksiz | NA |

| | | | | | | | |
|---|---|---|---|---|---|---|---|
| K. fedtschenkoi 2 | XX-0-TELA-2021.0039<br>RS 100 | tba | NA | NA | NA | NA | ex BG Rothschild |
| K. fedtschenkoi 3 | Zürich 99 5778 /0 =HtH 30791<br>W.Rauh M1323 | tba | W. Rauh | 11.1959 | Madagascar | Toliara, Fort Dauphin | This collection is recorded as Dracaena sp. In the BG Heidelberg accession books, and the number could be a mis-spelling of Rauh M1325, which is given as K. rosei in the accession books; received from BG Utrecht 16.10.2003 |
| K. aff. fedtschenkoi | München 2021/0736<br>Lyon 060320<br>RS 961 | tba | Richaud | 1994 | Madagascar | Tsivory | NA |
| K. spathulata var. garambiensis 2 | RS 521 | tba | Ikeda | NA | NA | NA | Steinhardt |
| K. gastonis-bonnieri | XX-0-TELA-2021.0042<br>RS 878 | tba | NA | NA | NA | NA | NA |
| K. deficiens 1 | München 2021/1712<br>PalmengartenFrankfurt 6-22817-1<br>RMY 219 | tba | Ralph Mangelsdorff | NA | Yemen | ca. 10 km Westlich Madinat ash Shirq 14°38'7.14"N 43°52'7.94"E ca.1500 m | NA |
| K. aff. glaucescens 1 | XX-0-TELA-2021.0043<br>RS 912 | tba | NA | NA | NA | NA | NA |
| K. x auriculata 2 | KE-0-B-0241074<br>Pfennig 1079 /3 | tba | Pfennig | NA | Kenya | NA | NA |
| K. aff. glaucescens 2 | UG-0-B-0241174<br>Pfennig 1078 | tba | Pfennig | NA | Uganda | NA | NA |
| K. glaucescens 1 | München 2021/0714<br>Lyon 210048 | tba | NA | NA | Ethiopia | NA | NA |
| K. deficiens var. deficiens | XX-0-B-0248474 | tba | NA | NA | NA | NA | NA |
| K. aff. glaucescens 4 | Berlin 024-86-74-30<br>Bally,P.R.O. & Melville,R. 16003 | tba | P.R.O.Bally & R.Melville | 18.1.1970 (?) | Somalia | Mait Pass | NA |
| K. aff. glaucescens 3 | XX-0-TELA-2021.0044<br>RS 672<br>CR7105 | tba | NA | NA | NA | NA | ex Jean-André Audissou |
| K. stearnii 2 | München 2021/1487<br>Mainz 198775260 | tba | NA | NA | Rwanda | NA | NA |
| K. gracilipes | München 2021/1491<br>Zürich 150182/0 | tba | NA | NA | Madagascar | NA | Anonymous 723 (ex BG Berlin) |
| K. grandiflora | München 1993/2942 | tba | NA | NA | NA | NA | NA |
| K. guignardii var. schistosepala | XX-0-TELA-2020.0463<br>RS 212 | tba | Richaud | 1994 | Madagascar | Ifaty, north of Tulear | NA |
| K. hildebrandtii var. glabra | Heidelberg 141045 | tba | NA | NA | Madagascar | NA | Donor: R. Hebding, 1995-03-00 |
| K. hirta | G.F. Smith 1141 | PRU 128254 | G.F. Smith | 1987 | SouthAfrica | KwaZulu-Natal | NA |
| K. humifica | Heidelberg 104872<br>RMM 451 | tba | R.D.Mangelsdorff | 3.-4.1998 | Madagascar | Bemaraha, ca. 25-30 km E Antsalova, growing together with Aloe (Lomatophyllum) antsingyense on top limestone dome amidst dense semideciduous forest, shady | Donor: R. D. Mangelsdorff, 1998-05-04 |
| K. humilis | G.F. Smith 1081 | PRU 125935 | G.F. Smith | 2001 | Mozambique | Nampula, province, near Nampula | NA |
| K. hypseloleuce | RS 274 | tba | NA | NA | NA | NA | NA |
| K. inaurata | München 2021/1545<br>Zürich 99 8451/0 | tba | NA | NA | Madagascar | NA | NA |
| K. spathulata var. spathulata | XX-0-TELA-2021.0066<br>RS 284 | tba | NA | NA | India | NA | ex Julie Finn |
| K. spathulata var. garambiensis 1 | TW-0-JENA-7784804 | tba | Arndt & Bopp | 17/09/2016 | Taiwan | Kenting Nationalpark, Karstwald östl. des Botanischen Gartens, verkarsteter Felsen, Korallenkalk, 220m (L: 120°49'22"E / B: 21°57'44"N) | NA |
| K. integrifolia var. bitteri | Zürich 15 0253 /0 | tba | Röösli & Hoffmann | 11.1992 | Madagascar | "Itremo" / "Col d'Itremo" | received from Röösli 10.11.2015 |
| K. jongmansii subsp. jongmansii | München 2021/1546<br>Heidelberg 140662<br>W. Rauh 22398 | tba | W. Rauh | 09/05/1969 | NA | Fianarantsoa Province, Amoron'i Mania Region, 3 km S Ambatofinandrahana, Matten, 1600 m, coordinates available | Donor: W. Rauh, 1969-09-00 |
| K. aff. laciniata | XX-0-TELA-2021.0048<br>RS 595 | tba | NA | NA | NA | NA | Tanzania? |
| K. laciniata | VN-0-LYJB-190117<br>München 2021/1495=2021/0744<br>Zürich 13 0121/0 | tba | Haager & Rybkovz | NA | Vietnam | Pongour Waterfalls | ex Jardin Exotique |
| K. laetivirens 2 | München 2021/1547<br>Zürich 99 8453/0<br>Descoings 28234 | tba | Descoings | 1994 | Madagascar | Toliara | NA |
| K. laetivirens 1 | Heidelberg 141072 | tba | NA | NA | Madagascar | SW-MG | Donor: H. Nothelfer, 1995-00-00 |
| K. lateritia 3 | München 2021/1496<br>Zürich 89 1751/0 | tba | Pocs | 1988 | NA | NA | ex BG Vacratot |
| K. lateritia 2 | XX-0-TELA-2021.0049<br>RS 197 | tba | NA | NA | NA | NA | NA |
| K. lateritia 1 | Berlin 196-08-88-20<br>Pfennig,H. 1740 | tba | H. Pfennig | 01/06/1988 | Kenya | Mombasa, 2 km östlich von Mackinnon Road | NA |
| K. lateritia var. prostrata | SO-0-B-2230183 | tba | Bally & Melville | 18/01/1973 | Somalia | Mait-Pass. | Samm.-Notes: Ost-Usambara 1979, leg Raadts & Nuernberg; die Pflanze wurde in Ostafrika gesammelt und von Prof. Nuernberg in seinem Gewächshaus kultiviert. Nach seinem Tod nahm der BG Hamburg die Pflanze in Kultur. Von dort sind Stecklinge nach Berlin geschickt worden. |
| K. lateritia var. zimbabwensis | XX-0-TELA-2021.0050<br>RS 091 | tba | NA | NA | NA | NA | ex Galin Radkov |
| K. lateritia x crenata | München 2021/1548<br>Berlin 146027920 | tba | NA | NA | Kenya | NA | NA |
| K. latisepala | München 2021/1549<br>Zürich 88 1142/0 | tba | Lavranos | NA | Malawi | NA | NA |
| K. laxiflora | RS 817 | tba | Aldo Torrebruno & Joël Jacq | NA | Madagascar | Toliara | NA |
| K. laxiflora var. subpeltata | XX-0-TELA-2021.0051<br>RS 807 | tba | Richaud | NA | NA | NA | Tsimbazaza? |
| K. laxiflora var. violacea | XX-0-TELA-2021.0052<br>RS 623 | tba | NA | NA | NA | NA | NA |
| K. leblanciae | G.F. Smith 1142 | PRU 128255 | G.F. Smith | 1993 | SouthAfrica | KwaZulu-Natal province, near border with Mozambique | NA |
| K. linearifolia 2 | München 2021/1500<br>Heidelberg 141029 | tba | NA | NA | Madagascar | S-MG, Cap Ste. Marie | Donor: unknown, 1969-00-00 |
| K. linearifolia 1 | München 2021/1499<br>Düsseldorf 201700055 | tba | NA | NA | Madagascar | Cape Sainte Marie | NA |
| K. lobata | Berlin 084-24-83-50<br>BG Zimbabwe 144/83 | tba | NA | NA | Zimbabwe | Zimbabwe: Südafrika, Zimbabwe, ohne Standortangaben | NA |
| K. lokarana | XX-0-B-2303774 | tba | NA | NA | NA | NA | NA |
| K. longiflora 2 | XX-0-TELA-2021.0053<br>RS 079 | tba | NA | NA | NA | NA | NA |
| K. longiflora 1 | Berlin 145-93-74-80 | tba | NA | NA | SouthAfrica | NA | provided by: Antwerpen, 12.3.1968; |
| K. lubangensis | G.F. Smith 1104 | PRU 127839 | G.F. Smith | 1999 | Angola | Namibe province. Bibala road from Humpata, below Leba Pass. | NA |
| K. luciae | G.F. Smith 1144 | PRU 128257 | G.F. Smith | 1981 | SouthAfrica | Mpumalanga province, near eManzana | This is the white to yellowish white flowered form. Material of the pink-flowered form not available. |
| K. manginii | München 2021/1502 | tba | M. Lehmann | NA | SouthAfrica(?) | NA | ex Bochum |
| K. marmorata 1 | München 2021/1514<br>Zürich 99 8417/0<br>Bally 12339 | tba | Bally | 1961 | Kenya | Rift Valley Prov. | NA |
| K. marmorata 2 | Berlin 197-02-05-10 | tba | U. Katz | NA | Ethiopia | Harar, Mt. Achim | provided by: Bochum, Botanischer Garten der Ruhr-Universität Bochum; |
| K. marmorata var. somaliensis | XX-0-B-0470707 | tba | NA | NA | NA | NA | NA |

| | | | | | | | |
|---|---|---|---|---|---|---|---|
| K. marnieriana | München 2021/1506<br>Heidelberg 140666<br>Rauh 69755 | tba | NA | NA | Madagascar | Plateau | Les Cèdres |
| K. maromokotrensis | XX-0-TELA-2021.0055<br>RS 619 | tba | Rebmann | NA | Madagascar | sud du massif de l'Analamera, Maromokotra | ex P. Richaud |
| K. millotii | München 2021/1507<br>Heidelberg 140583<br>Rauh 74635a | tba | W. Rauh | 1959 | Madagascar | Toliara Province, Ambovombe | Donor: W. Rauh, 1959-00-00 |
| K. miniata 2 | München 2021/0732<br>Lyon 210099<br>Röö/Hoffm. 78/98 | tba | Röösli & Hoffmann | NA | NA | NA | NA |
| K. miniata 1 | München 2021/1508<br>Zürich 94 1521/0<br>Lavranos 9562 | tba | Lavranos | 1972 | Madagascar | Fianarantsoa | NA |
| K. miniata 3 | Berlin 231-13-90-30 | tba | NA | NA | Madagascar | | provided by: SSZ 86 3639; sonstige Herkunft: Leuenberger |
| K. mitejea | Berlin 230-63-74-80<br>Pfennig,H. 1460 | tba | H. Pfennig | NA | Kenya | NA | NA |
| K. neglecta | G.F. Smith 1145 | PRU 128258 | G.F. Smith | 1995 | SouthAfrica | KwaZulu-Natal province, eastern parts of the province | NA |
| K. aff. olivacea | München 2021/1551<br>Zürich 99 9312 / 1 | tba | Billensteiner | 1991 | Myanmar | NA | NA |
| K. orgyalis | München 2021/1510<br>Heidelberg 140582<br>Rauh 7408 | tba | W. Rauh | 15/10/1961 | Madagascar | Faritany [Province], Toliara [Tuléar], Région Atsimo-Andrefana, Fivondronana [Sous-préfecture], Ankazoabo, inhabited place [Andriambe / Andriabe], coordinates available, Trockenwald | Donor: W. Rauh, 1961-10-21 |
| K. paniculata | G.F. Smith & E. Figueiredo 44 | PRU 123653 | G.F. Smith & E. Figueiredo | 1986 | SouthAfrica | North West province, Hartebeespoortdam. | NA |
| K. pareikiana | SER23-034 | tba | David-Paul Klein, Ronen Shtein, S | 24/10/2023 | Madagascar | Near Ambondrofe on small Tsingys beside the road | NA |
| K. peltata var. mandrakensis | SER23-042 | tba | Seraina E. Rodewald, David-Paul | 27/10/2023 | Madagascar | near Mandraka Forest along the road (-21.252167, 47.892658) | |
| K. peltigera | München 2021/1512<br>Zürich 99 8455/0 | tba | Richaud | NA | Madagascar | NA | ex Descoings |
| K. perrieri | XX-0-TELA-2020.0469<br>RS 909<br>Lyon 060321 | tba | Richaud | 1994 | Madagascar | South Madagascar | NA |
| K. petitiana | München 1979/0598 | tba | NA | NA | NA | NA | NA |
| K. aff. petitiana | München 2021/0734<br>Lyon 210074 | tba | NA | NA | NA | NA | Originally from HBG |
| K. pinnata 3 | XX-0-TELA-2021.0058<br>RS 420 | tba | NA | NA | NA | NA | Ikeda s.n. Tsimbazaza |
| K. pinnata 1 | AHM 311 | tba | A.M. Havinga & Iharivolana | 17/02/2022 | Madagascar | province de Diego-Suarez/Antsirana, sous-préfecture de Ambilobe, localité de Manambato. Forêt de Sorata. Coord. Précises (WGS 84): 13.70644°S, 49.45333°E. Altitude 917m. Près d'un chemin dans une zone déforestée. Substrat: roches mouillées, crevasses remplis d'humus, habitat naturel probablement dans un environnement perturbé ensoleillé, en présence de Cynorkis sp.. Herbacée, 30cm de haut, non fertile; rebord des f.rouges | Récolté avec D.P. Klein, échantillon photographié in situ. |
| K. pinnata 2 | München 2021/1513<br>Mainz 198575390 | tba | NA | NA | Rwanda | NA | NA |
| K. prasina | G.F. Smith 1082 | PRU 125936 | G.F. Smith | 2001 | Mozambique | NA | without precise locality |
| K. prittwitzii 1 | XX-0-TELA-2021.0060<br>RS 985<br>ISI 2013-24 | tba | NA | NA | NA | NA | NA |
| K. prittwitzii 3 | München 2021/1482<br>Zürich 99 9304/0 | tba | NA | NA | NA | NA | ex cult. BG Darmstadt |
| K. prittwitzii 2 | München 2021/1516<br>Berlin 024397430 | tba | NA | NA | Kenya | NA | NA |
| K. aff. prittwitzii | XX-0-B-0470607 | tba | NA | NA | NA | NA | NA |
| K. prolifera | XX-0-B-0554874 | tba | NA | NA | NA | NA | NA |
| K. pubescens 1 | XX-0-B-0234774 | tba | NA | NA | NA | NA | NA |
| K. pubescens 2 | München 2021/1517<br>Heidelberg 141008<br>D. Supthut 86213 | tba | D.Supthut | 16/10/1986 | Madagascar | 25 km S of Ambositra, on granite rocks | Donor: Sukkulenten-Sammlung Zürich, 1990-08-00 |
| K. pumila | RM-1-HEID-140680<br>Rauh 8094 | tba | NA | NA | Madagascar | NA | NA |
| K. quartiniana | München 2021/0728<br>Lyon 210081<br>RS 088 | tba | NA | NA | NA | NA | NA |
| K. rebmannii | München 2021/1553<br>Zürich 19 0141/0 | tba | Solichon | NA | NA | NA | ex BG Meise |
| K. Zebedi | XX-0-TELA-2020.0471<br>Descoings 422<br>RS 258 | tba | NA | NA | NA | NA | NA |
| K. rhombopilosa 2 | N/0174 | tba | NA | NA | NA | NA | NA |
| K. rhombopilosa 1 | Heidelberg 140327 | tba | E. Mengarelli | NA | Madagascar | NA | Donor: E. Mengarelli, 1987-10-00 |
| K. robusta | München 2021/1556<br>Düsseldorf 201700003 = YE-0-DUSS-5966 | tba | NA | NA | Yemen | N-YE, Hanus-G, Menacha Al Hoteib, 2400m | NA |
| K. rolandi-bonapartei | München 2021/1714<br>Palmengarten Frankfurt 21-31767-4<br>RMM 244 | tba | Ralph Mangelsdorff | NA | Madagascar | Antsiranana, Montagne d'Ambre | NA |
| K. rotundifolia | XX-0-TELA-2021.0064<br>RS 938<br>WY 1092 | tba | NA | NA | Yemen | southwestern Yemen | NA |
| K. stearnii 3 | München 2021/1519<br>Zürich 99 3107/0<br>Lavranos & Smith 431 | tba | Lavranos & Smith | NA | Yemen | Socotra | ex Les Cèdres |
| K. aff. crenata 1 | München 2006/1136 | tba | NA | NA | NA | NA | NA |
| K. stearnii 1 | München 2021/1522<br>Heidelberg 145140<br>Rauh 3003 | tba | W. Rauh | 10/07/1961 | SouthAfrica | Province Gauteng, inhabited place City of Tschwane Metropolitan Municipality, Hammanskraal, coordinates available | Donor: BG Berlin-Dahlem, 2014-06-23 |
| K. rubella | München 2021/0730<br>Lyon 210082<br>RS 612 | tba | Descoings | NA | NA | NA | ex Richaud |
| K. sanctula | München 2021/1557<br>Zürich 99 8457/0<br>Descoings 28180 | tba | Descoings | 1994 | Madagascar | Amboasary, Taolagnaro region | NA |
| K. scandens | München no number | tba | NA | NA | NA | NA | public greenhouse behind glass next to Welwitschia |
| K. schimperiana | München 2021/0726<br>Lyon 210084<br>WB 663 | tba | NA | NA | Ethiopia | South of Abaro. Portuguese Bridge | NA |
| K. schizophylla | XX-0-B-0236574 | tba | NA | NA | NA | NA | NA |
| K. serrata 2 | XX-0-TELA-2021.0065<br>RS 628 | tba | NA | NA | NA | NA | NA |
| K. serrata 1 | München 2021/1524<br>Heidelberg 141019 | tba | NA | NA | Madagascar | Fianarantsoa Province, Manara | Donor: unknown, 1997-05-00 |
| K. sexangularis | München 2021/1525<br>Heidelberg 106009 | tba | NA | NA | SouthAfrica | NA | NA |
| K. sexangularis var. intermedia | XX-0-TELA-2004.0005<br>RS 808 | tba | NA | NA | NA | NA | ex BG Vienna, Austria |
| K. sp. 3 | Paris 1975 V477-43379 | tba | NA | NA | Madagascar | NA | Tsimbazaza |

| | | | | | | | |
|---|---|---|---|---|---|---|---|
| K. sp. 1 | XX-0-TELA-1997.0051 | tba | NA | NA | NA | NA | NA |
| K. aff. faustii | XX-0-TELA-1997.0045<br>RS 130 | tba | NA | NA | NA | NA | ex BG Jibou, Zimbabwe |
| K. streptantha | München 2021/0733<br>Lyon 210086<br>ISI 2013-26 | tba | NA | NA | NA | NA | NA |
| K. suarezensis 2 | XX-0-TELA-2021.0068<br>RS 425 | tba | NA | NA | NA | NA | NA |
| K. suarezensis 1 | XX-0-TELA-2021.0069<br>RS 750<br>ISI 90-68 | tba | NA | NA | NA | NA | NA |
| K. synsepala | München 2021/1528<br>Heidelberg 145141<br>W.Rauh 10757 | tba | W. Rauh | 17/08/1963 | Madagascar | Fianarantsoa Province, 24 km E Ambatofinandrahana, coordinates available, Granitfelsen | Donor: BG Berlin-Dahlem, 2014-06-23 |
| K. tenuiflora | XX-0-TELA-2020.0467 == XX-0-TELA-2021.0070<br>RS 266<br>Descoings 28300 | tba | NA | NA | NA | NA | Tsimbazaza |
| K. tetraphylla | München 2021/1530<br>Heidelberg 140331<br>Rauh 22473 | tba | W. Rauh | 21/05/1969 | Madagascar | Faritany [Province] Fianarantsoa, Région Amoron'i Mania, Fivondronana [Sous-préfecture] Ambatofinandrahana, inhabited place Ampanetoana [?] (24 km E Ambatofinandrahana), coordinates available, schattige Felsspalten | Donor: W. Rauh, 1969-06-00 |
| K. cf. thyrsiflora | München 2021/1531<br>Heidelberg 142184<br>Rauh 58002 | tba | W. Rauh | 01/03/1982 | SouthAfrica | Western Cape, Stellenbosch | Donor: W. Rauh, 1982-00-00 |
| K. tomentosa | München 2021/1532<br>Heidelberg 140352<br>Rauh M 722a | tba | W. Rauh | 02/10/1959 | Madagascar | Fianarantsoa Province, Vatovavy-Fitovinany (region), Ifanadiana (district), coordinates available, Bergwald von Ranomafana | Donor: W. Rauh, 1959-10-12 |
| K. torrejacqii | XX-0-TELA-2020.0468<br>RS 580 | tba | A. Torrebruno & J. Jacq | NA | Madagascar | Namorona river valley | NA |
| K. delagoensis | XX-0-TELA-2021.0071<br>RS 424<br>HBG 789 | tba | Charles Swingle & Henri Humbert | 1928 | NA | NA | NA |
| K. usambarensis | TZ-0-LYJB-200058<br>München 2021/0743 | tba | NA | NA | Tanzania | Province de Tanga | NA |
| K. variifolia var. seyrigii | XX-0-B-0236174 | tba | NA | NA | NA | NA | NA |
| K. variifolia var. variifolia | XX-0-TELA-2021.0063<br>RS 426 | tba | Richaud | NA | Madagascar | Behara | NA |
| K. lateritia var. lateritia | G.F. Smith 1119 | PRU 127922 | G.F. Smith | 1997 | Zimbabwe | Central. Masvingo and Lake Mutirikwe | NA |
| K. viguieri | München 2021/1715<br>Palmengarten Frankfurt 13-27140-1<br>RMM 4103 | tba | Ralph Mangelsdorff | NA | Madagascar | Tulear | NA |
| K. waldheimii | RS 244 | tba | Richaud | 1994 | Madagascar | Ibity | NA |
| K. fedtschenkoi 1 | München 2021/1559<br>Heidelberg 141025 | tba | NA | NA | Madagascar | NA | Donor: Les Cèdres, 1990-06-00 |
| K. waterbergensis | G.F. Smith & E. Figueiredo 90 | PRU 127835 | G.F. Smith & E. Figueiredo | 1999 | SouthAfrica | Limpopo province, northwest of Rhenosterpoort, Jan Trichardt Pass | NA |
| K. crenata | XX-0-TELA-2005.0012<br>RS 043 | tba | NA | NA | NA | NA | NA |
| K. sp. 2 | XX-0-B-2300574 | tba | NA | NA | NA | NA | NA |
| K. winteri | G.F. Smith & E. Figueiredo 50 | PRU 125606 | G.F. Smith & E. Figueiredo | 1999 | SouthAfrica | Limpopo province, Wolkberg, Thabakgolo Escarpment, Sedibeng sa Lebese Mountain | NA |
| K. x flaurantia | XX-0-TELA-2020.0456<br>RS 863<br>Descoings 28305 | tba | NA | NA | Madagascar | Amboasary | NA |
| K. x rechingeri | München N/3127 | tba | NA | NA | NA | NA | NA |
| K. yemensis | Berlin 233-05-86-30 | tba | NA | 3.1981 | Yemen | Sanaa (Prov.), Wadi Dhar bei Sanaa, Sandstein, 2200m. | NA |
| Tylecodon buchholzianus | Heidelberg 142180<br>Rauh 11540 | tba | W. Rauh | 27/09/1963 | SouthAfrica | Northern Cape, Annisfontein | Donor: W. Rauh, 1963-00-00 |
| Tylecodon torulosus | Heidelberg 102659<br>Rauh 30183 | tba | NA | NA | SouthAfrica | Northern Cape Province, Richtersveld, Karrachabfoort | Donor: J. J. Lavranos, 1971-00-00 |

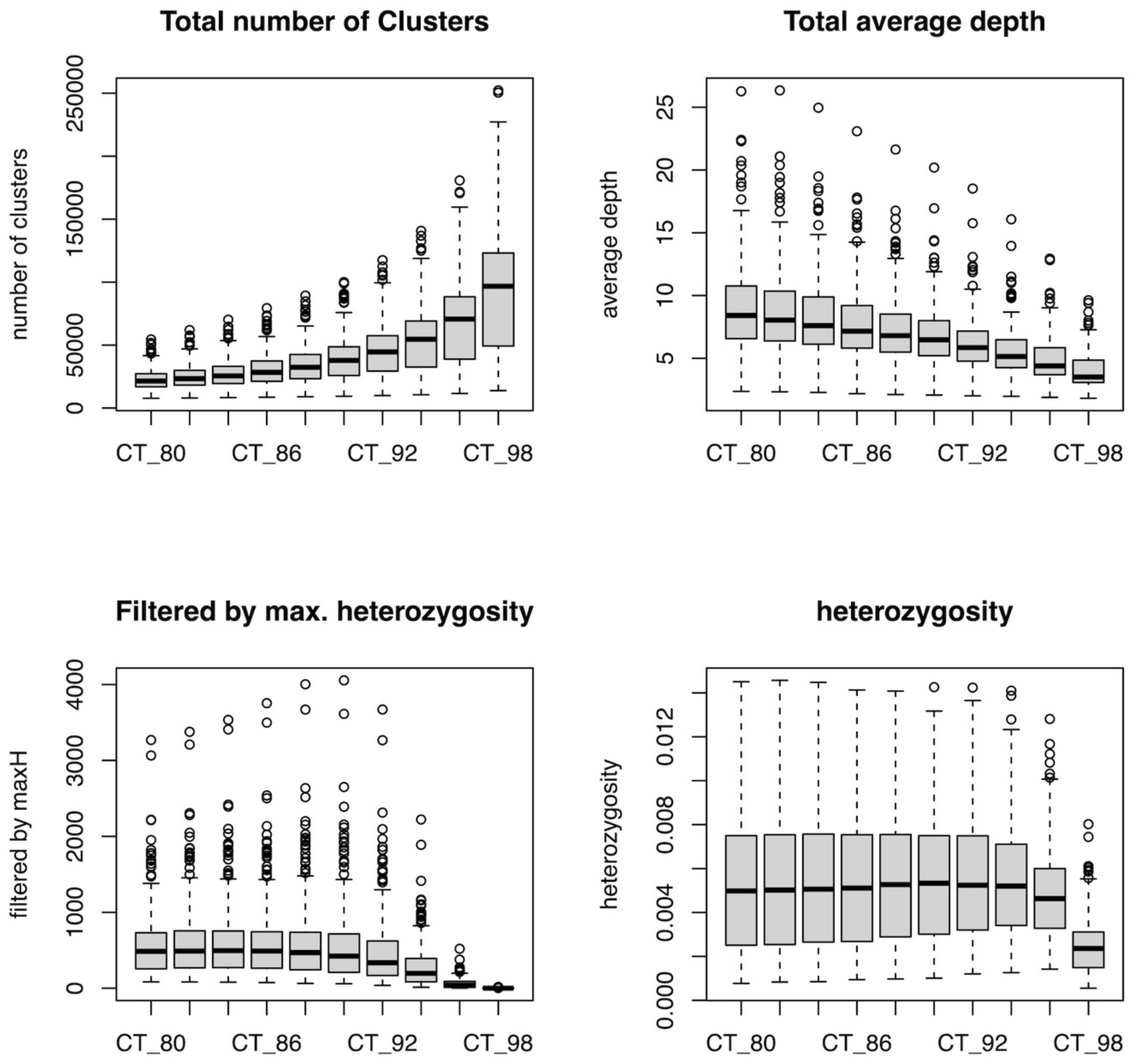


Supplementary Figure S1a. Comparison of within-sample clustering thresholds for four metrics that were evaluated for the assembly with the Ipyrad pipeline. The total number of clusters, total average depth, loci filtered by maximum heterozygosity, and heterozygosity per sample are shown across within-sample clustering thresholds of 80–98% similarity between reads.

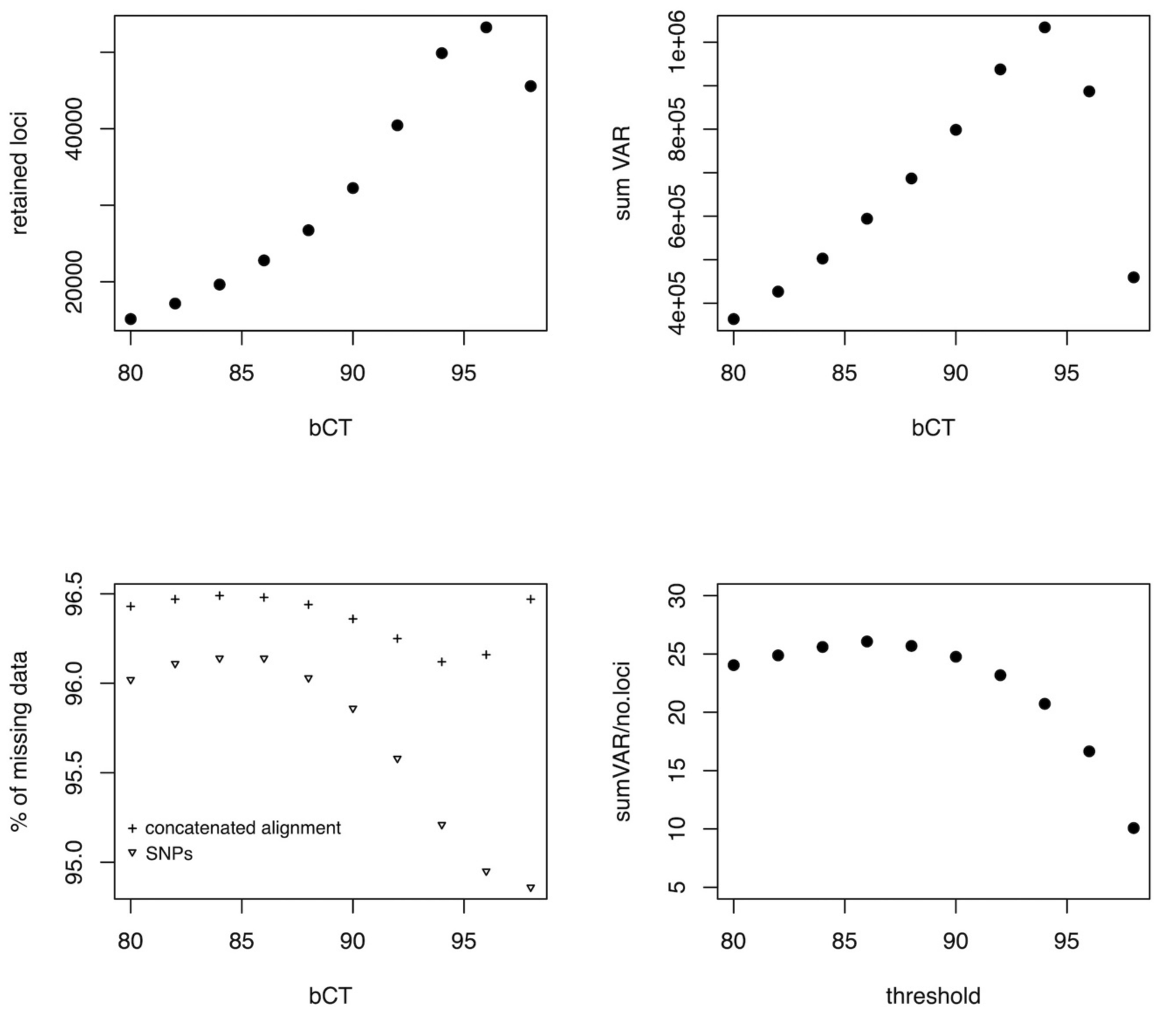


Supplementary Figure S1b. Comparison of between-sample clustering thresholds for four metrics that were evaluated for the assembly with the Ipyrad pipeline. The total number of retained loci, the total number of variable sites, the percentage of missing data and the average number of variable sites per locus are compared across between-sample clustering thresholds of 80–98% similarity between within-sample consensus sequence clusters.

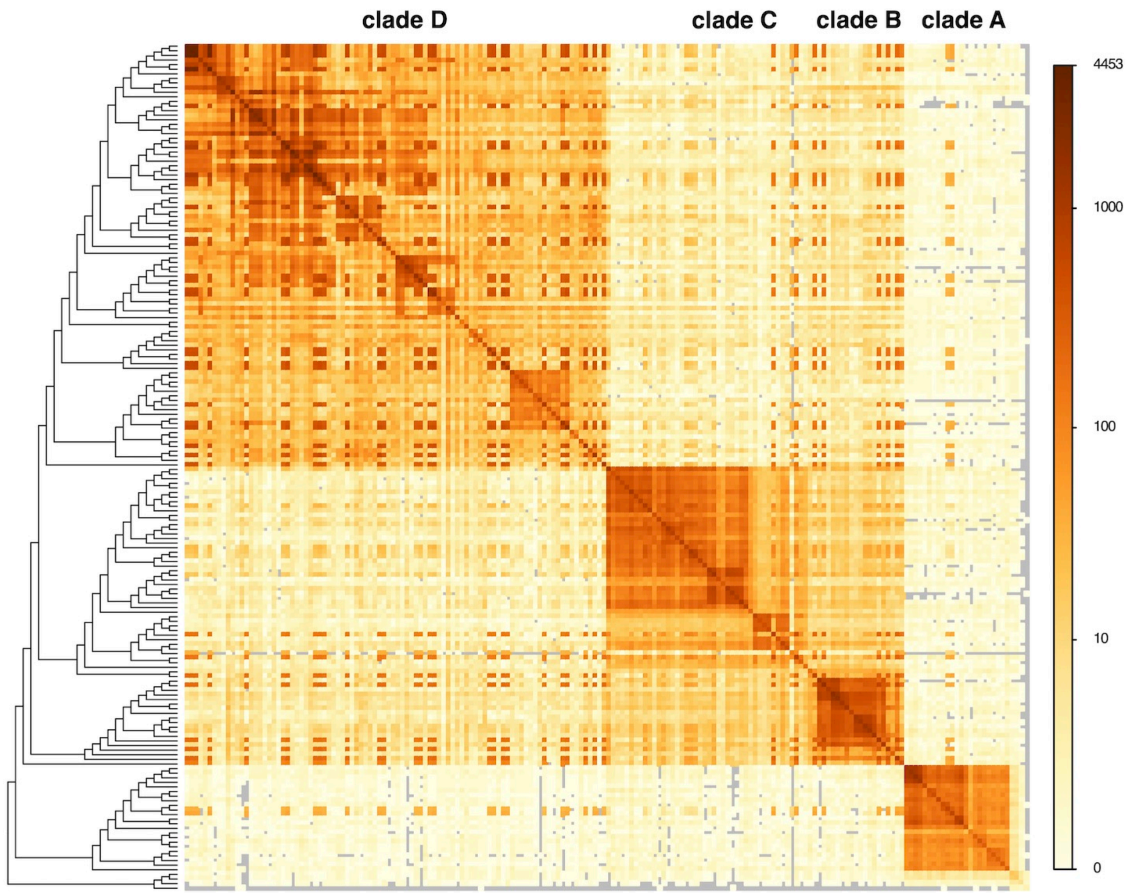


Supplementary Figure S2. Number of ddRADseq loci shared between samples. Rows and columns represent individual samples. They are ordered according to their placement in the concatenated tree, shown on the left. The diagonal entries show the number of loci recovered for each sample and off-diagonal entries represent the number of shared loci between the samples. The colour coding is log-scaled with darker colours representing more shared loci between samples.

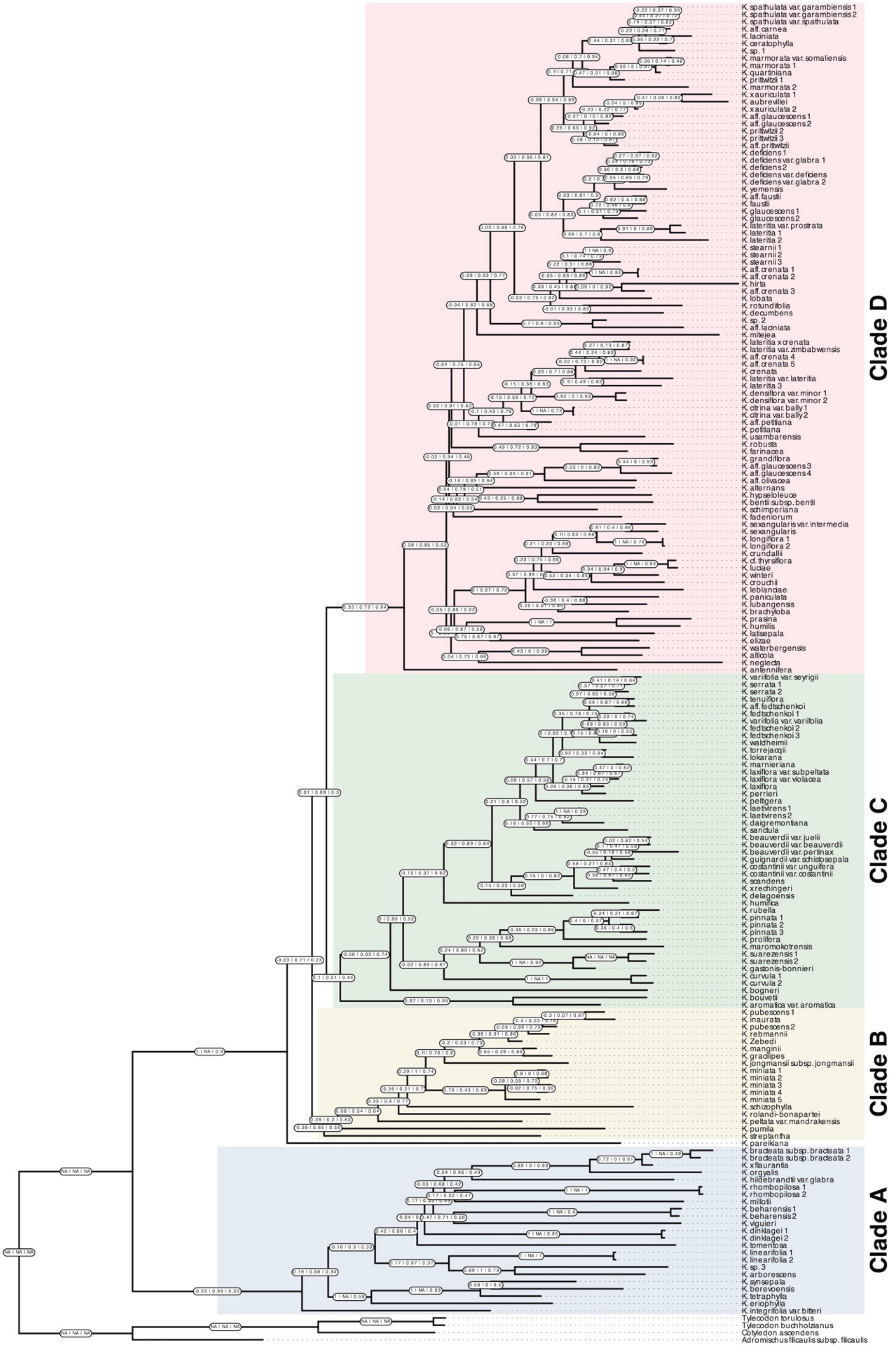


Supplementary Figure S3. Quartet sampling scores on maximum likelihood phylogeny of *Kalanchoe s.l.* inferred from the concatenated alignment of ddRADseq loci (concatenated tree). Quartet sampling scores (Quartet concordance (QC), quartet discordance (QD) and quartet informativeness (QI)) are shown for each branch separated by a vertical line.

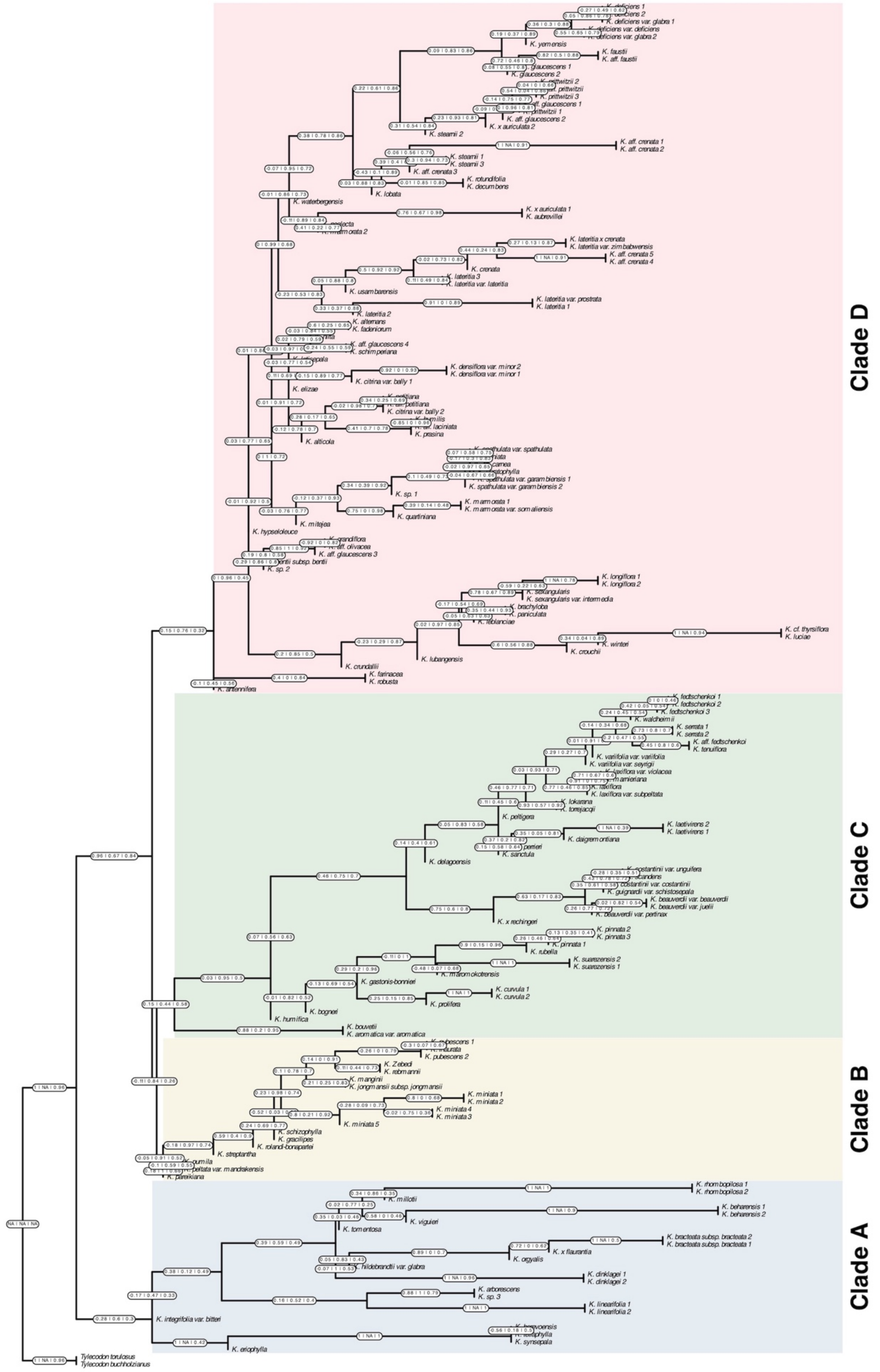


Supplementary Figure S4a. Quartet sampling scores on ddRADseq phylogeny of *Kalanchoe s.l.* inferred with a two-step coalescent approach (coalescent tree). Quartet sampling scores (Quartet concordance (QC), quartet discordance (QD) and quartet informativeness (QI)) are shown for each branch separated by a vertical line.

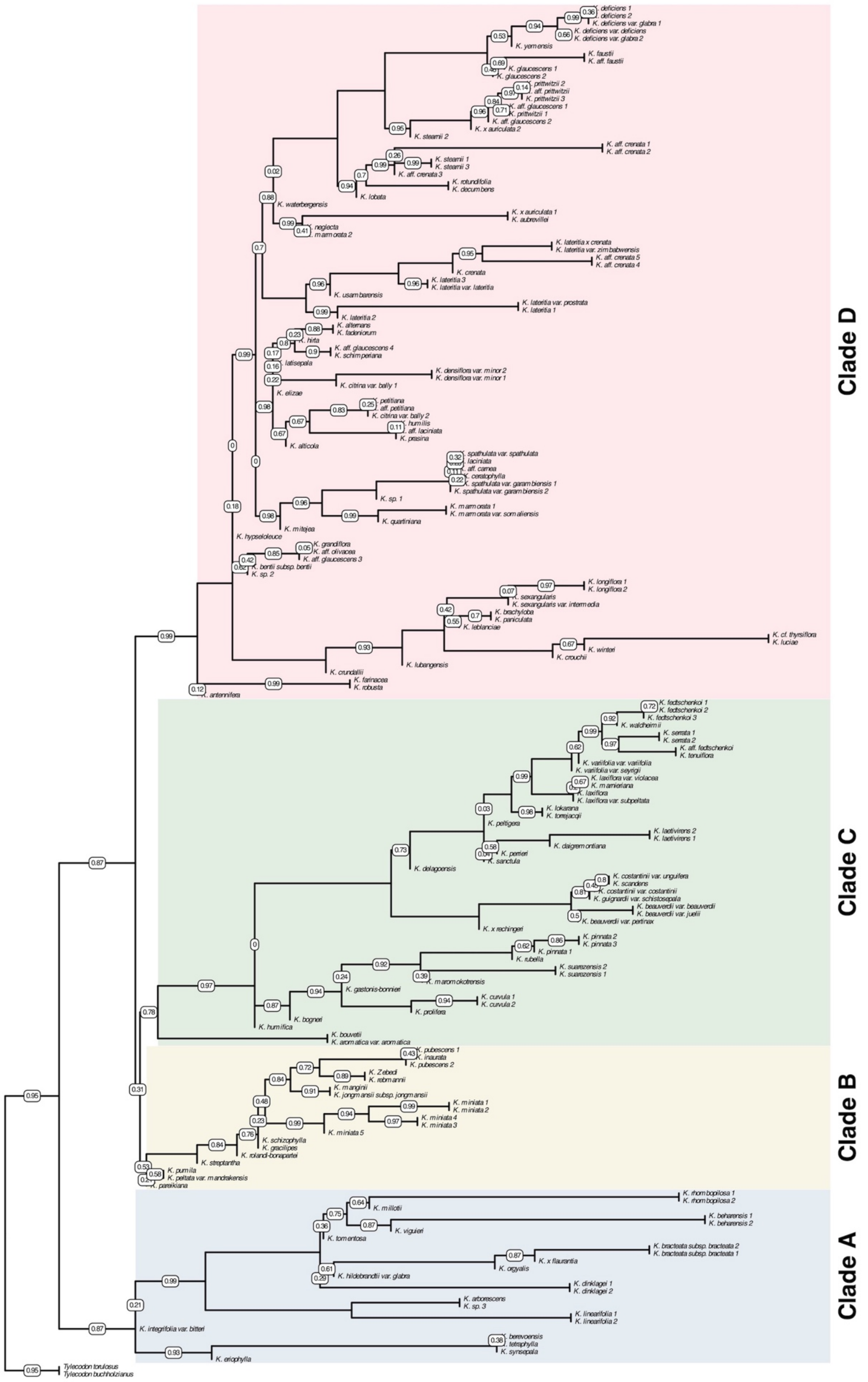


Supplementary Figure S4b. Local posterior probability support values on ddRADseq phylogeny of *Kalanchoe s.l.* inferred with a two-step coalescent approach (coalescent tree). local posterior probability support values are shown only for nodes that do not receive maximum support

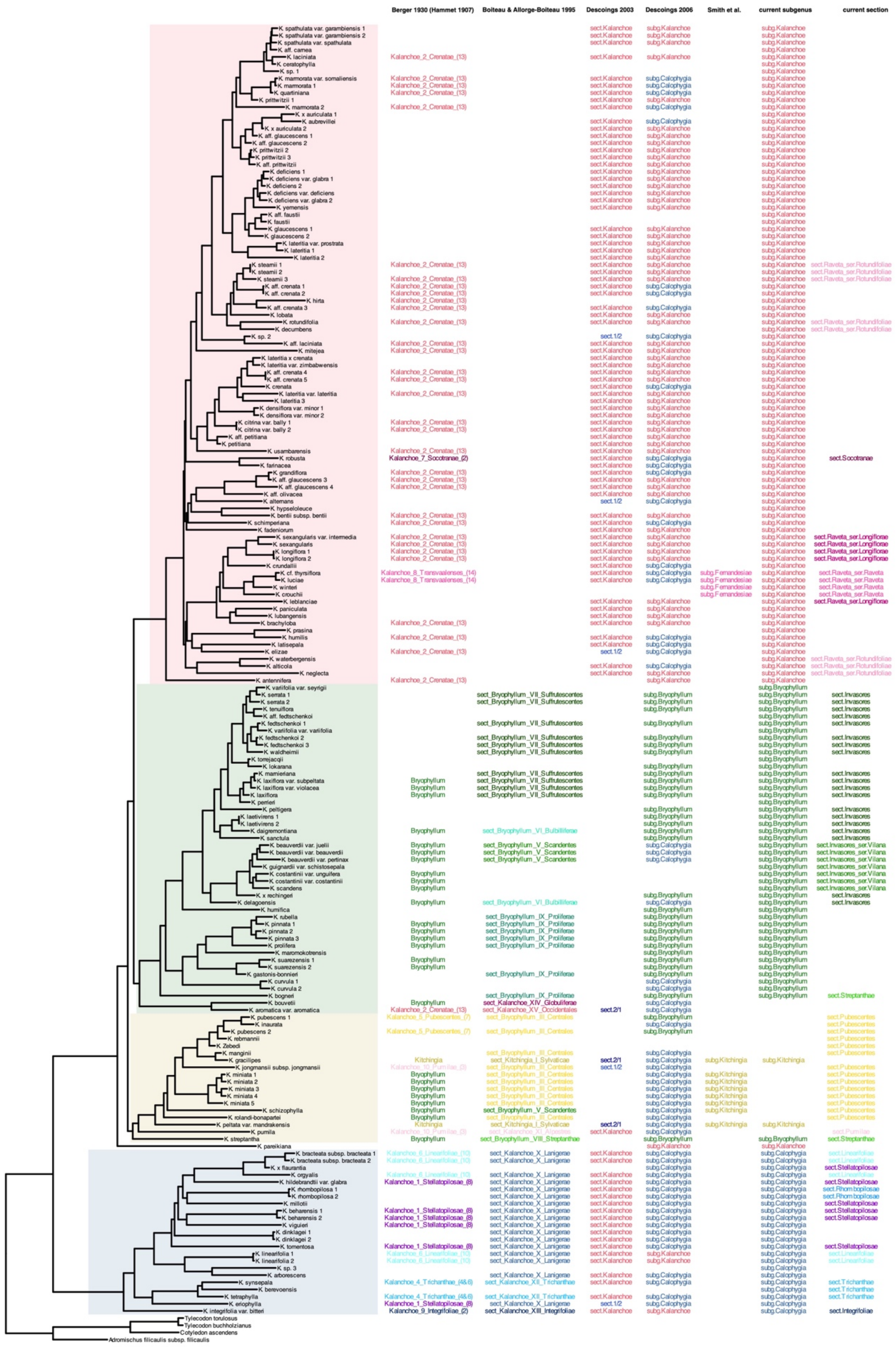

Supplementary Figure S5. Partial taxonomic history of *Kalanchoe s.l.*: Comparison of selected taxonomic classifications from 1930 to the present in relation to the concatenated tree. Berger (1930) treated *Kalanchoe s.s.*, *Bryophyllum*, and *Kitchingia* as separate genera and divided *Kalanchoe* into 10 unranked units largely based on the numbered groups of

Hamet (1907; 1908). Boiteau and Allorge-Boiteau (1995) recognised three sections in *Kalanchoe s.l.*: *Kitchingia*, *Bryophyllum*, and *Kalanchoe*. They furthermore split the genus into 15 informal groups. Descoings (2003) recognised two sections only in *Kalanchoe s.l.*: *Kalanchoe* and *Bryophyllum*. Affinities of species intermediate between the two sections were so indicated. Descoings (2006) recognised three subgenera in *Kalanchoe*: *Kalanchoe*, *Bryophyllum*, and *Calophygia*, with those species that he considered to be intermediate between *K.* subg. *Kalanchoe* and *K.* subg. *Bryophyllum* contained in *K.* subg. *Calophygia*. Smith and Figueiredo (2018) published a combination for *Kitchingia* at the rank of subgenus and Smith (2021) published the name *K.* subg. *Fernandesiae*. The circumscription of these two taxa has since been superseded. The last two columns represent the most recent taxonomic treatment at the ranks of subgenus and section (including series), largely representing the work of one of us (GFS) and colleagues since 2018. Smith (2023*g*; 2024*a*) provide comprehensive reviews of the infrageneric taxonomic history of *Kalanchoe s.l.*.
